# Hedgehog Signalling Modulates Glial Proteostasis and Lifespan

**DOI:** 10.1101/2020.02.05.935585

**Authors:** Andrew Rallis, Juan A. Navarro, Mathias Rass, Amélie Hu, Serge Birman, Stephan Schneuwly, Pascal P. Thérond

**Affiliations:** Université Côte d’Azur, CNRS, Inserm, iBV, France; Department of Developmental Biology, Institute of Zoology, Universitaetsstr. 31, University of Regensburg, 93040 Regensburg, Germany; Genes Circuits Rhythms and Neuropathology, Brain Plasticity Unit, ESPCI Paris, CNRS, Labex MemoLife, PSL Research University, 10 rue Vauquelin, 75005 Paris, France

## Abstract

The conserved Hedgehog (Hh) signaling pathway has a well-established role in animal development, however its function during adulthood remains largely unknown. Here, we investigated whether the Hh signaling pathway is active during adult life in *Drosophila melanogaster* and uncovered a protective function for Hh signaling in coordinating correct proteostasis in glial cells. Adult-specific depletion of Hh reduces lifespan, locomotor activity and dopaminergic neuron integrity. Conversely, increased expression of Hh extends lifespan and improves fitness. Moreover, Hh pathway activation in glia rescues the lifespan and age-associated defects of *hedgehog* (*hh*) mutants. At the molecular level, the Hh pathway regulates downstream chaperones, principally *hsp40* and *hsp68*, whose overexpression in glial cells rescues the shortened lifespan and proteostasis defects of *hh* mutants. Finally, we demonstrate the protective ability of Hh signalling in a *Drosophila* Alzheimer’s disease model expressing human Amyloid Beta (Aβ1-42) in the glia. Overall, we propose that Hh signalling is requisite for lifespan determination and correct proteostasis in glial cells and may have potential in ameliorating a wide range of degenerative diseases.

## INTRODUCTION

The Hedgehog (Hh) signalling pathway is an evolutionarily conserved module that organizes animal patterning and is essential for determining cell identity and the final body plan (Briscoe and Thérond., 2013). Hh and components of its pathway were initially discovered in *Drosophila* and highly conserved members are present in the vast majority of metazoa including in mammals, which possess three Hh paralogs, the most well studied being Sonic Hedgehog (Shh; Ingham, 2018). In human, abnormally sustained activation of the Shh pathway in the adult is critical in the initiation and growth of many tumors (Beachy et al., 2004; Curran, 2018). Although the developmental facets of Hh signalling have been well established, the role of Hh signalling in adult homeostasis has been difficult to evaluate due to the vital function of the Hh pathway during development.

Emerging lines of evidence suggest that Shh has adult specific functions in the nervous system (Petrova and Joyner, 2014). For instance, in mice, Shh regulates the proliferation of neural stem cells throughout adult life (Lai et al., 2003; Palma et al., 2005; Ahn and Joyner, 2005; Ferent et al., 2014; Yao et al., 2016). Shh signalling also has been demonstrated to act as a modulator of brain plasticity and nerve regeneration (Yao et al., 2016). Furthermore, Shh signaling is elevated in different CNS injury models, such as stroke, spinal cord injury and stab wound (Jin et al., 2015; Amankulor et al., 2009; Bambakidis et al., 2010). In line with the above, treatment with Shh or Shh agonists has shown to improve neurological function and promote neuroregeneration (Chechneva et al., 2014; Huang et al., 2013; Bambakidis et al., 2012; Thomas et al., 2018) whereas depletion of Shh signalling components exacerbated the neuropathology following cerebral ischemia and nerve injury (Ji et al., 2012; Martinez et al., 2015).

There is accumulating evidence that Shh signalling utilises a neuron-glia communication mode to coordinate homeostasis in the post-natal brain. Under physiological conditions, Shh is predominantly released by neurons (Gonzalez Reyes et al., 2012; Eitan et al., 2016; Okuda et al., 2016) and glial cells have been shown, *in vitro*, to be highly responsive to Shh signalling (Ugbode et al., 2017; Okuda et al., 2016). Furthermore, Shh signaling is active in *vivo* in the adult mouse forebrain (Garcia et al., 2010) in which signaling is activated in a discrete population of glial cells that respond to Shh secreted from neurons (Garcia et al., 2010, Farmer et al., 2016). Importantly, activating Shh signaling in glial cells has been demonstrated to be neuroprotective. For example, *in vitro*, Shh-stimulated mouse cortical astrocytes protect co-cultured neurons from kainite (KA)-induced cell death (Ugbode et al., 2017). Furthermore, in neurodegeneration mouse models induced by intraperitoneal injection of KA, or surgical brain injury, Shh signaling in glia regulates proliferation of astrocytes and microglia (Sirko et al., 2013; Pitter et al, 2014).

Due to its role as a neuroprotective factor, it is hypothesised that Shh may be implicated in alleviating age-related neurodegenerative disease (Zhang et al., 2014; Yao et al., 2017). For instance in cultured rat hippocampal neurons, Shh signalling confers protection against neurotoxins such as the Amyloid beta peptide (Aβ 1−42; Yao et al., 2017). However the mechanism by which Shh exerts its protective function in a live organism has not been explored. In Alzheimer’s disease, accumulation of Aβ 1−42 has been demonstated to precede the formation of amyloid plaques, and the deposition of amyloid plaques correlates with an increase in age-related cognitive dysfunction (D’Andrea et al., 2001; Rodrigue et al, 2009). Aβ 1−42 has a high propensity to aggregate and there is a large body of evidence that chaperones prevent deleterious misfolding and aggregation of Amyloid (Evans et al, 2006; Magrané et al, 2004). In fact, the Hsp70 chaperone and its partner Hsp40 have been demonstrated to act synergistically to inhibit the early stages of Aβ 1−42 aggregation *in vitro* (Evans et al, 2006). Moreover in rat primary neuronal cultures seeded with Aβ 1−42, *hsp70* overexpression improves cell viability (Magrané et al, 2004). Based on previous studies is possible that Shh prevents accumulation of Aβ 1−42 aggregates, however this has yet to be evaluated.

Here we utilize *Drosophila melanogaster* to evaluate the function of the Hh pathway at the physiological and cellular level during adult life. This model enables us to address the tissue-specific functional role of the Hh signaling pathway in the adult brain. We identify a new role for Hh signalling in adult life span and glial proteostasis and uncover the mechanism by which Hh prevents accumulation of protein aggregates. We show that increasing Hh levels promotes lifespan extension, whereas depletion of Hh activity during adult life reduces lifespan. We further demonstrate that specific activation of the Hh pathway in adult glia potently rescues the shortened lifespan, locomotor and dopaminergic neuron defects observed in *hedgehog* (*hh*) mutants. We found that proteostasis in glial cells is affected in absence of Hh activity and identified two highly conserved chaperones *hsp40* and *hsp68* (a member of the Hsp70 protein family) that are regulated by the Hh pathway. Their overexpression potently rescues proteostasis defects in the glia of *hh* mutant animals and rescues *hh* mutant reduced lifespan. As proof of concept we also demonstrate that activation of the Hh pathway and one of its downstream chaperone targets in the glia is able to rescue the deleterious effects of expression of human Amyloid Beta (Aβ 1-42) in an *in vivo* Alzheimer’s disease model. The work described here makes a significant contribution to our understanding of the role of the Hh morphogen during adulthood: it identifies for the first time a protective role for Hh on glia proteostasis, and dissects its mechanism of action, showing that Hh operates through the regulation of highly conserved chaperones.

## RESULTS

### The Hh signaling pathway is present in glial cells of the adult brain

Here, we address the role of Hh signaling in the adult brain of Drosophila melanogaster in which it has not yet been studied. To do so, we first investigated the pattern of expression of Hh signalling components, in particular the Smoothened (Smo) serpentine protein and the transcription factor Ci/Gli which is able to activate the expression of target genes (Fig. 1a-d). For this, we utilized the Crispr/Cas9 genome editing and generated a new Ci-Trojan Gal4 line which represents the full Ci expression pattern and, together with a second independent system (Ci-LexA line), found that Ci is predominantly expressed in glial cells in the adult fly brain (Fig. 1a, b and Fig. S1a). Immunofluorescent analysis using antibodies against Smo and Ci was also performed and revealed a strong localization of both components in glia (Fig. 1c-d). RNAi-depletion of Smo and Ci in glia reduced immunoreactivity for both proteins, supporting specificity of the stainings (Fig. S1b-e). These findings are consistent with a recent single cell transcriptome analysis of the *Drosophila* and human adult brain identifying Smo and Ci/Gli expression in glia (Davie et al., 2018, Lake et al, 2018).

**Figure 1.**
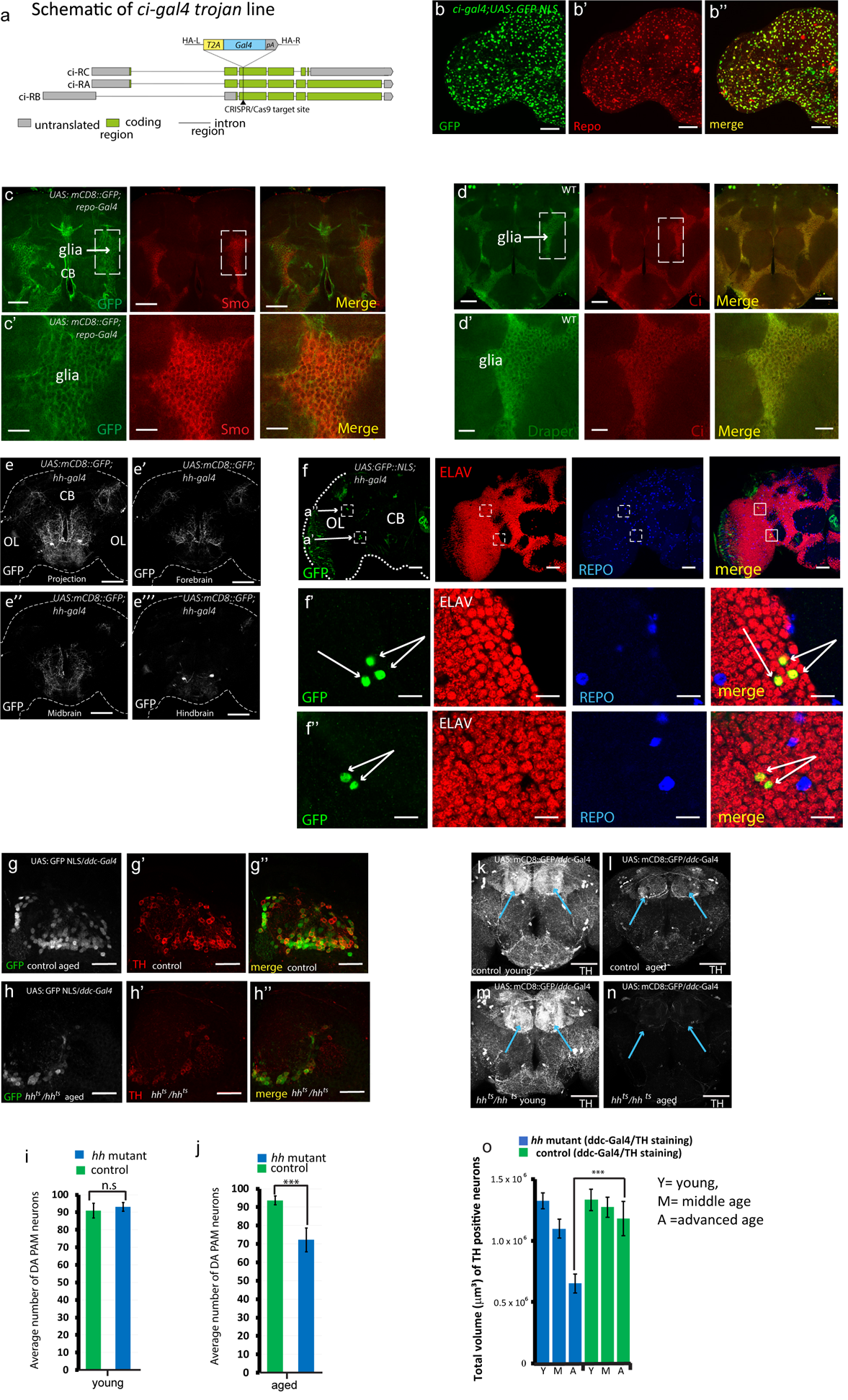
Location of Hh signalling components in the *Drosophila* adult brain and DA neuron loss in *hh* mutants. (a) Schematic representation of the Ci-T2A-Gal4 trojan line. (b-b”) Cells positive for ci – T2A-Gal4 (trojan line) expressing GFP NLS (green channel – b) are immunostained with the glial marker Repo (red channel-b’). The merged image (b’’) reveals that almost all the cells expressing Ci are glial cells. Scale bars: 50 μm.(c-d) Endogenous Smo and Ci are localised in the glial cells. Anterior 1μM section of a whole mount adult brain in which glial cells are specifically labelled with (c) mCD8::GFP (green channel) or with (d) anti-Draper (a glial receptor, green) and immunostained for Smo or Ci (red channel). Note that Smo and Ci staining are present in glial cells (dashed boxes), scale bars: 50 μm. (c’-d’) Magnification of the region of the glia indicated by the dashed boxes. Scale Bars: 20 μm. (e-f) *hh*-expressing cells in the adult brain have neuronal identity. Projection pattern for Hh expressing cells (e-e’’’). Cells positive for *hh-*Gal4 express membrane marker mCD8::GFP, (e-e’’’) images of 1μm sections for the forebrain, midbrain and hindbrain. (f) Expression of a nuclear GFP in *hh*-expressing cells of adult brain are positive for Elav (red channel), but not Repo (blue channel). Scale Bar: 50μm. (f’-f’’) 5x magnification of the boxed regions shown in (f). The merge between *hh*-expressing cells (green) and anti-Elav immunostaining (red channel) is shown in the right panel. *hh*-expressing cells are indicated by white arrows. Scale bars: 5 μm. (Images consist of a 1 μm section). (g-h’’) Representative confocal images of the PAM cluster cell bodies of control (g-g’’) and *hh* mutant (h-h’’) of aged adult fly brains, in which DA neurons are labelled with a nuclear GFP marker (GFP NLS in grey) and with anti-TH immunostaining (in red). Scale bars:10μm. (i-j) Quantifications of DA neuron number in the PAM cluster of young and aged flies. (k-n) *hh* mutant and control brains with DA neurons immunostained with TH (in grey) in young (k, m) and aged flies (l, n). Arrows label the PAM DA neurons. (o) Quantifications of DA neuron volume in control and *hh* mutant adult fly brains with TH immunostaining, in young (Y), middle (M) and aged flies (A). For DA neuron number statistics the two tailed T-test was used and for DA neuron volume quantification two ways analysis of variance (ANOVA) test was used to compare the specified data points for *hh* mutant and control samples (***, p < 0.001; **, p < 0.01 and *, p < 0.05).

Knowing that the Hh pathway is present in adult glial cells, we further examined the pattern of expression of Hh in the adult brain. Using two different *hh*-Gal4 reporter lines, inserted at two different positions in the *hh* genomic locus, we found that *hh* is expressed in the adult brain in approximately 10-15 neurons per hemisphere that are positive for the pan-neuronal marker Elav, and negative for the glia-specific marker Repo (Fig. 1e-f and S1f-g). Using a third reporter line for *hh*-expressing cells – the *hh* dsRed enhancer trap (Akimoto et al., 2005) - we identified glutamatergic and GABAergic neurons that are positive for the *hh*-expressing cells (Fig. S1h-j). This shows that both subtypes of neurons produce Hh, in agreement with recent single cell transcriptome analysis of the *Drosophila* adult brain (Croset et al., 2018; Davie et al., 2018) and with an equivalent study on Shh in the adult human brain (Lake et al, 2018).

### Hh signalling is needed to maintain Dopaminergic neuron viability in the adult brain

To investigate the functional relevance of Hh signalling in adult, we specifically depleted Hh in the entire organism using a *hh* temperature sensitive allele (*hh^ts^*). This is a null allele for *hh* at non-permissive temperature (29°C) and, after 24h at this temperature, no more Hh staining or activity can be detected in homozygote mutant animals (Ranieri et al., 2012; Palm et al., 2010). To avoid any patterning defect due to Hh developmental function, *hh*^ts^ flies were raised at permissive temperature (18°C) and transferred to non-permissive temperature upon eclosure (Methods). Indeed, these flies do not exhibit any noticeable patterning defect (Strigini et., 1997). With this set up, we investigated whether Hh is required to maintain neuronal homeostasis in the adult brain. To explore this, we focused on dopaminergic (DA) neurons known to be highly susceptible to brain homeostasis defects, and whose cell number, neuronal projection and secretion of dopamine can be measured and quantified accurately (Friggi-Grelin et al., 2003; Navarro et al., 2014).

Nine DA clusters are present in the Drosophila adult brain, among which the protocerebral anterior medial – PAM – cluster, that is part of the circuit regulating climbing activity (Riemensperger et al., 2013). Strikingly, using anti-Tyrosine hydroxylase (TH, a functional marker for DA neurons) immunostaining (Friggi-Grelin et al., 2003), we observed a reduced number of DA neurons in the PAM cluster of aged *hh* mutant adult brains (Fig. 1g-h and j) whereas no change in DA neuron number was detected in young *hh* mutant flies (Fig. 1i, S2f-g). These defects suggest that overall DA neuron volume in the adult brain might be affected in absence of Hh. Indeed, we found that, during adulthood, *hh* mutant brains globally showed an abnormally low level of TH staining (Fig. 1k-o, Table S1) as well as an overall reduction of the mRNA levels of the neuronal specific form of TH (nTH) and a decrease in brain dopamine levels (Fig. 3n-o, compare green and blue bars). In an alternative approach, we quantified DA neuron volume network by labelling their cell membrane with the mCD8-GFP reporter transgene and found similar results (Fig. S2a-e). To do so, we used the Ddc-Gal4 driver which targets a subset of DA neurons including the entire PAM cluster (Riemensperger et al., 2013). Indeed, Ddc - DOPA decarboxylase - is the enzyme that catalyzes the transformation of L-Dopa into dopamine. Taken together these results show that Hh is necessary to maintain DA homeostasis in adult brain, which is illustrated with the substantial reduction of DA neuron functional integrity observed in *hh* mutant flies.

### Adult expression of Hh is necessary for correct lifespan determination

We next decided to investigate whether the *hh* mutant flies exhibit any survival or behavioral deficits, since defects in the dopaminergic neuron network have been demonstrated to have a detrimental effect upon lifespan and locomotor activity in both mammals and invertebrates (Thanos et al., 2016; Liao et al., 2017; Feany and Bender, 2000). Strikingly, we found that adult *hh^ts^* flies shifted to 29°C exhibit a drastically shortened lifespan (80 - 85% decrease in the median lifespan when compared to control flies), both females and males (Fig. 2a, S3c and Table S2). When maintained at 25°C, but not at 18°C, the *hh^ts^* mutant flies also exhibited a strongly decreased lifespan (Fig. S3k, Table S2).

**Figure 2.**
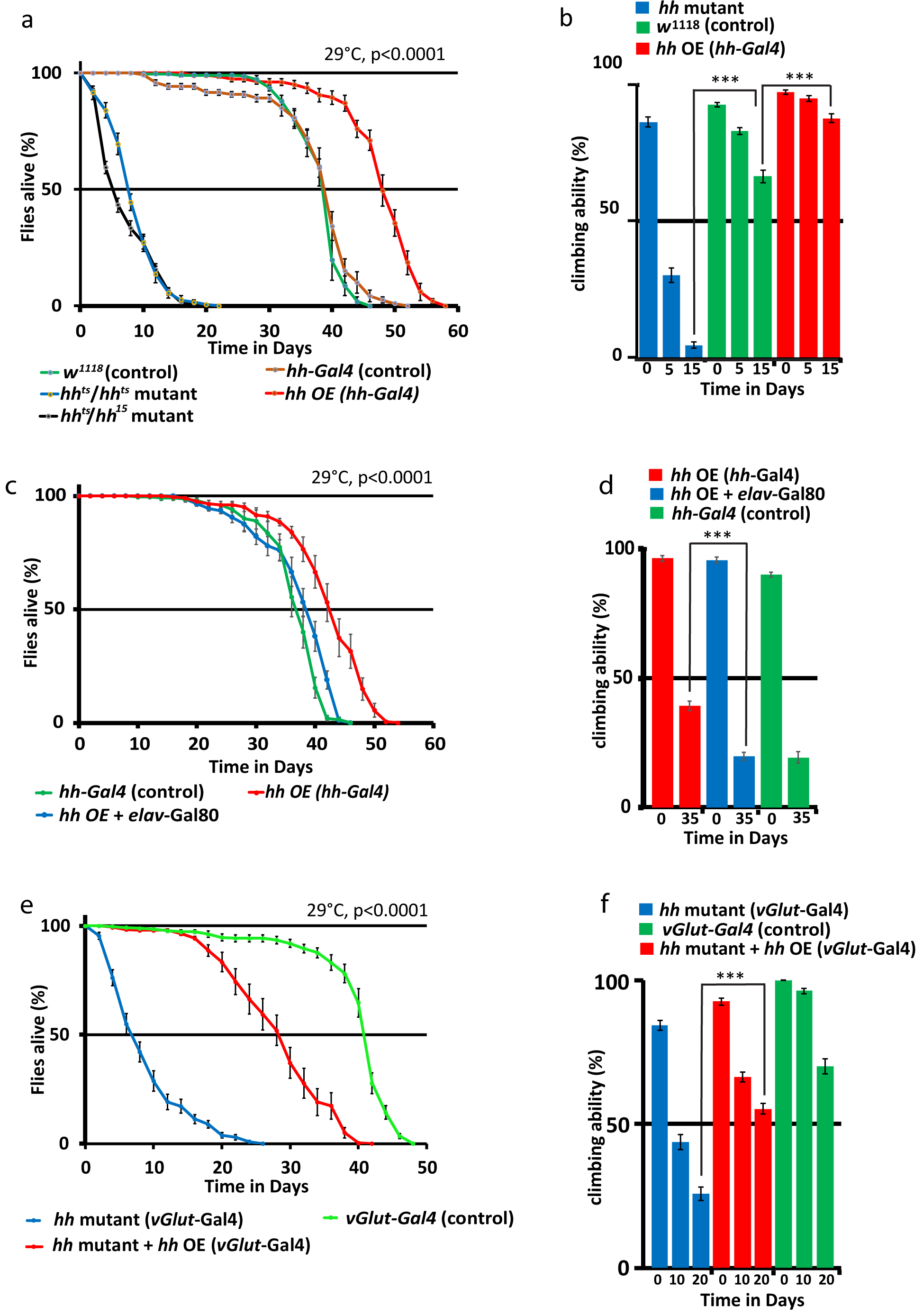
Hh signalling affects adult lifespan determination and fitness. (a) Overexpression of *hh* in *hh*-expressing cells - *hh* OE (*hh* Gal4) - extends survival rate whereas *hh* depletion drastically reduces it. Both combinations of *hh* mutant alleles (*hh*^ts^/*hh*^ts^ and *hh*^ts^/*hh*^15^) decrease the survival rate. (b) *hh* overexpression in adult flies shows improved negative geotaxis ability during adult life compared to control flies whereas aged hhts/hhts mutant flies exhibit strong climbing defects. (c-d) The *hh*-dependent lifespan extension phenotype and improved fitness - *hh* OE (hh Gal4) - is abolished upon repression of *hh* expression in neurons - *hh* OE (elav-Gal80) - but not in glia (Table S2). (e-f) Expression of *hh* in glutamatergic neurons of *hh* mutant - *hh* mutant + *hh* OE (vGlut-Gal4) - efficiently rescues the short-lived phenotype of *hh* mutants and the impaired locomotion phenotype. *p* values refer to the degree of difference between the *hh* mutant and control/rescue samples as well as *hh* OE and equivalent control at median survival. Detailed genotypes, % of changes in median survival and climbing activity, as well as corresponding *p* values can be found for all figures in tables S2-3 and S5-6. For all lifespan studies the log rank test was utilized, to compare median lifespans of two specified data sets, and for all climbing activity assays (Table S2-S3 and Table S5-S6), the two way ANOVA test was used to compare the specified data points (***, p < 0.001; **, p < 0.01 and *, p < 0.05).

To confirm that the loss of Hh is responsible for this defect, we analysed the heteroallelic combination of *hh^ts^* null flies with *hh^15^*, another strong allele of *hh*. The *hh^ts^/hh^15^* mutant combination reduced lifespan by a similar degree as *hh^ts^* (Fig. 2a and Table S2). In addition, *hh^ts^* adult mutant flies exhibited diminished climbing ability when compared to age-matched controls (Fig. 2b and Table S3). To further confirm that this effect is specifically due to *hh* loss of function, re-expressing *hh* exclusively into glutamatergic neurons in *hh^ts^* mutant flies potently rescued the median lifespan (red curve Fig. 2e), reaching 68% of the control one (green curve Fig. 2e), whereas expression in GABAergic neurons was sufficient to rescue median lifespan in *hh^ts^* mutant flies up to 41% (S3a and Table S2). The *hh^ts^* climbing defects were also significantly rescued even in advanced aged flies (Fig. 2f, S3b and Table S3). The individual UAS and Gal4 transgenes alone had no significant effect on survival rate compared to control flies (Fig. S3d-e and Table S2). These results suggest that Hh expression in the adult fly brain is essential for correct lifespan determination and locomotion activity.

### Increased level of Hh results in lifespan extension

Then we tested whether increasing Hh level at adult stage extends lifespan. When *hh* was <u>o</u>ver<u>e</u>xpressed (*hh* OE) in an adult specific manner using a driver under the control of the *hh* promoter (see Methods and Table S2), we observed a considerable expansion (21%) of the median lifespan accompanied with a significantly improved climbing ability (Fig. 2a, b and Table S2 and S3). Interestingly, the *hh* OE-dependent phenotypes were almost completely abolished upon inhibition of neuronal *hh* OE (using elav-Gal80 repressor of Gal4, see Methods) (Fig. 2c-d, Table S2 and 3). This effect was not observed upon expression of Gal80 in glial cells (using repo-Gal80; Table S2). Altogether, these data suggest that the *hh*-dependent lifespan extension and fitness relates to *hh* expression in adult neurons, and that potentially a sub-population of both GABAergic and glutamatergic *hh*-expressing neurons contribute to this process.

### Hh signalling in glial cells regulates lifespan determination

In order to confirm the function of the Hh signaling module in glia, we analysed the consequence of *smo* or *ci* depletion in these cells. Glia-specific RNAi for *smo* or *ci* (using repo-Gal4) led to a strong reduction in survival rates, with a decrease of 43% and 48% in the median lifespan respectively (Fig. 3a and Table S2). In contrast, *ci* and *smo* depletion in neurons (using *elav*-Gal4, Fig. 3b and Table S2) only had a minor effect on survival rates. Furthermore, expressing an activated form of Ci (*Ci^act^*; Wang et al., 1999) in glial cells at 29 or 25°C strongly rescued the *hh^ts^* mutant shortened median lifespan and defective climbing ability (Fig. 3c-d, S4a-a”, S3k and Tables S2 and S3). Additionally, overexpression of *Ci^act^* or an activated form of *Smo (Smo^act^,* Zhang et al, 2004) in glia was sufficient to extend median survival time by 13% and 16% respectively (Fig. 3e-f). In all these experiments, the presence of each individual UAS transgene had a negligible effect on survival time (Fig. S3f-j and Table S2).

**Figure 3.**
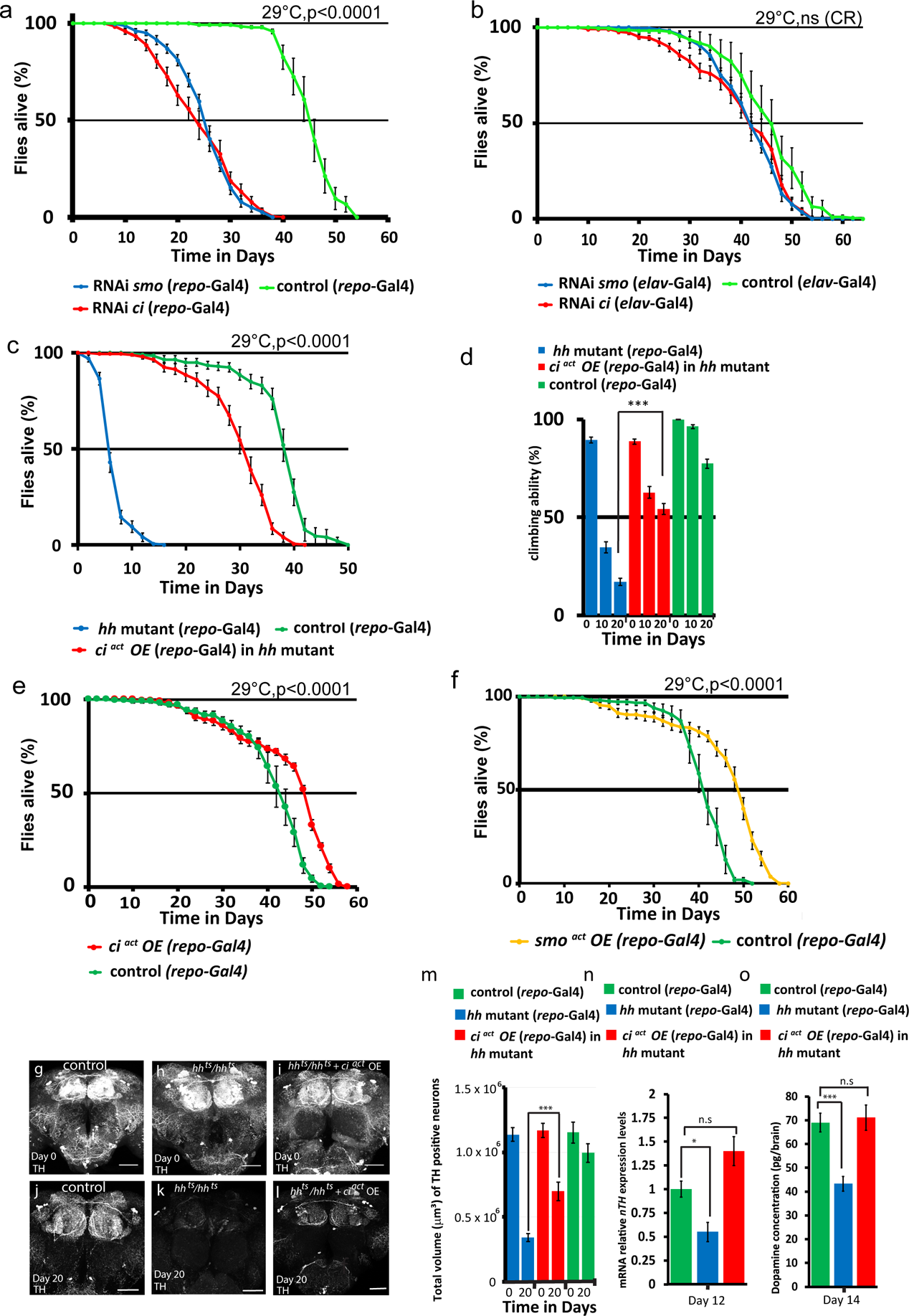
Hh signaling in glial cells determines Hh mediated lifespan and is protective. (a-b) Adult specific depletion of Smo (RNAi *smo*) or Ci (RNAi *ci*) in glia (*repo*-Gal4) but not in neurons (*elav*-Gal4) strongly reduces survival compared to the control. (b) Cox-regression^cr^ statistical analysis was used to compare control and RNAi lines and revealed no significant differences. (c-d) Expression of an activated form of Ci in the glia (*ci^act^* OE, *repo*-Gal4) rescues the short survival time of *hh* mutant, at 29°C well as (d) the impaired locomotion. (e-f) Overexpression of *ci^act^* (e) *or smo^act^* (f) in glial cells, extends lifespan (g-i) Images of newly eclosed day 0 fly adult brains with TH immunostaining, (g) control, (h) *hh* mutant, (i) *hh* mutant expressing *ci ^act^* in glia. (j-l) The expression of *ci ^act^* (l) in the glia of *hh* mutant flies, partially restores the volume of TH neurons exhibited in *hh* mutant flies (k) (quantified in m). Scale bars: 50μm, Z-projections of entire adult brain. (n-o) neuronal TH (nTH) mRNA levels and dopamine brain concentration are significantly reduced in *hh* mutant flies. Expression of *ci^act^* in the glia potently restores both (n) nTH mRNA expression and (o) brain dopamine to control levels. For lifespan studies the log rank test was utilized, and for climbing activity assays and DA neuron volume, the two-way ANOVA test was used (***, p < 0.001; **, p < 0.01 and *, p < 0.05, see Tables S1-S3 and S5-S6). Analysis of mRNA levels TH expression and Dopamine levels differences between samples were determined by a two-tailed t-test and one way ANOVA test respectively (***, P < 0.001; **, P < 0.01 and *, P < 0.05, see methods).

To confirm that the DA neuron defects we observed in *hh* mutant brains (described in Fig. 1) depends on the Hh signalling cascade in glial cells, we analysed whether expressing *Ci^act^* in glial cells of a *hh* adult mutant rescues the integrity of DA neurons. Indeed, expression of *Ci ^act^* in glia in a *hh* mutant background strongly recovered the DA neuron volume (Fig. 3g-m) as well as the low neuronal TH mRNA expression and reduced dopamine content (Fig. 3n-o). Overall our data demonstrates that transduction of Hh signal through Smo and Ci activity in glial cells is necessary and sufficient for *hh*-dependent lifespan and fitness. It also shows that Ci-dependent activity in the glia is essential in protecting DA neurons during adult life.

### Hh signaling regulates proteostasis in glial cells

To identify the Hh targets involved in neuroprotection and lifespan determination, we searched for potential protective genes downstream of Hh-dependent transcription in the adult brain. We performed a selective qRT-PCR screen for anti-oxidant defense molecules, autophagy components as well as chaperones (Fig. 4a and S5b-d), demonstrated to be implicated in lifespan regulation (Sun et al., 2012; Simonsen et al., 2008; Tower, 2011). As proof of principle, we examined well-established Hh specific targets regulated at the transcriptional level, such as *patched* (*ptc*, the canonical Hh receptor), *decapentaplegic* (*dpp*, of the TGFbeta family) and *knot/collier* (of the EBF transcription factor family) as well as other members of the Hh module which are post-translationally regulated (*ci* and *smo*; Fig. S5a). In agreement with reports analysing Hh targets during development (Biehs et al, 2010), we found that *ptc* was specifically downregulated in brains of flies depleted for adult Hh activity. Out of the protective factors tested in aged *hh* mutant adult brain samples, a distinct set of three chaperones, the heat shock proteins Hsp23, Hsp40 and Hsp68, displayed a significant mRNA level decrease, when compared to controls (Fig. 4b). Importantly, the expression of *Ci ^act^* in glial cells of *hh* mutants potently rescued the mRNA levels of *hsp23*, *hsp40* and *hsp68* (Fig. 4b), supporting the notion that *hsp* transcriptional downregulation is a consequence of the loss of Hh signaling. Furthermore, mRNA levels of both *hsp40* and *hsp68* are similar in aged-matched control flies at both 25°C and 29°C, whereas both chaperone mRNA levels are drastically elevated at 37°C heat shock conditions (Fig. 4c-d). This shows that lifespan condition run at 29°C exhibit no significant difference in regards to the mRNA levels of *hsp40* and *hsp68* compared to 25°C conditions.

**Figure 4.**
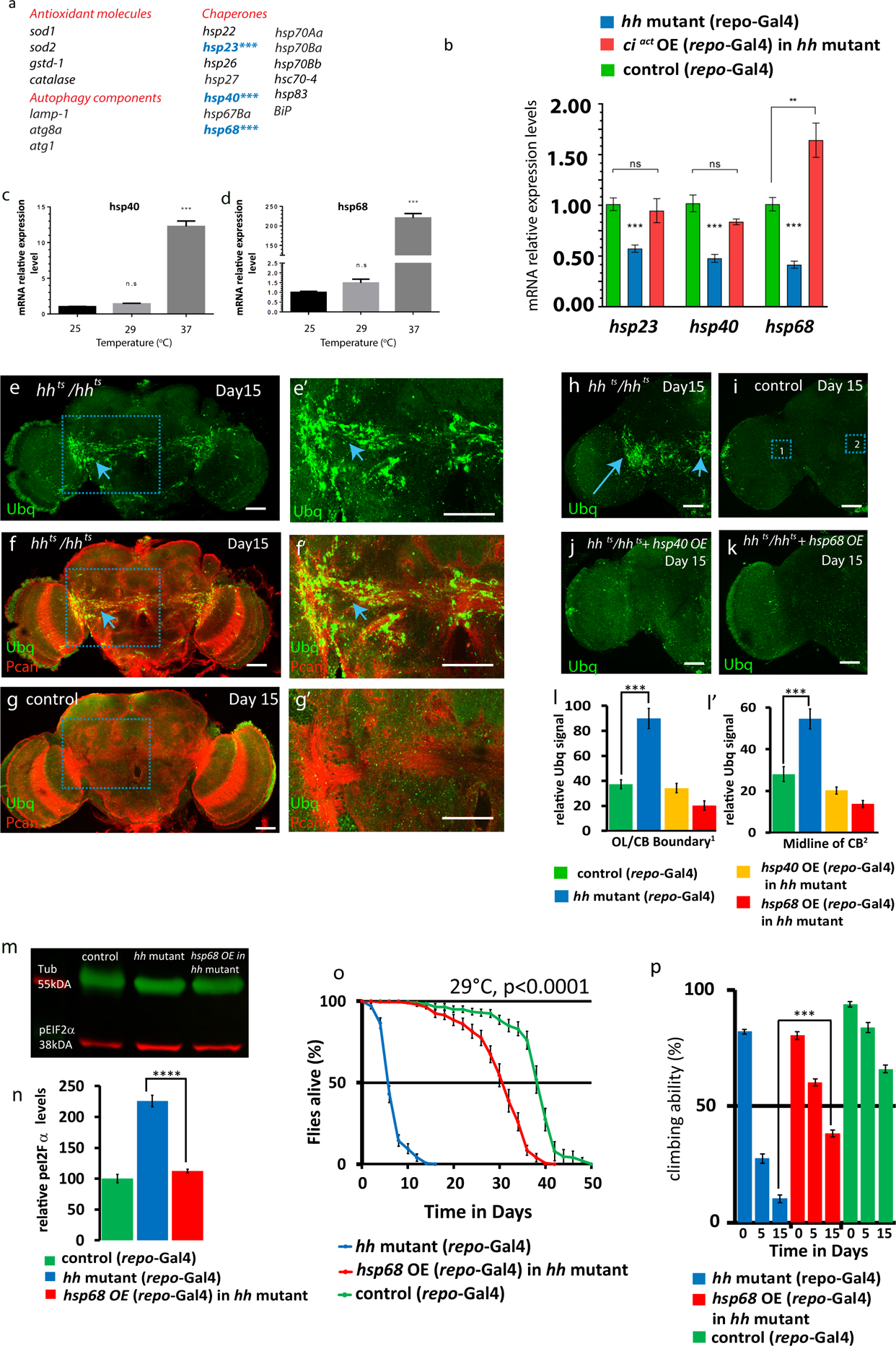
*hh* target chaperones maintain glial homeostasis, survival and fitness. (a) List of protective factors tested in a qRT-PCR screen. mRNA levels significantly decreased in adult *hh* mutant heads are highlighted in blue whereas mRNA levels of other chaperones and genes implicated in antioxidant defences and in lysosomal autophagy pathway (Fig. S4) show no significant difference. (b) The mRNA levels of *hsp23*, *hsp40* and *hsp68* chaperones are all significantly decreased in 10-day-old *hh* mutants (two-way t-test, p <0.001***) and restored to close to or above control levels in *hh* mutant flies expressing *ci^act^* in the glia. (c-d) Relative mRNA expression level of *hsp40* (c) and *hsp68* (d) at 25°C, 29°C and 37°C for day 12 control flies. No significant difference (One-way ANOVA, ns) was observed in mRNA levels of *hsp40* and *hsp68* between 25°C and 29°C. (e-g) Day 15 *hh* mutant and control brains immunostained for Ubiquitin (Ubq, green) and Perlecan (Pcan, red in f, g). Scale bars 50μm. (e’-g’) Zooms of squared areas labelled in e-g. Scale Bars 30μm, 1 μM section. Ubiquitin positive aggregates (blue arrows) are predominantly located in midbrain glial tracts immunostained with Pcan (f’). (h-k) Day 15 *hh* mutant (h) and control (i) brains and (j-k) *hh* mutant flies expressing *hsp40* (j) or *hsp68* (k) in glia, immunostained for anti-ubiquitin. Ubiquitin aggregates (blue arrows) are observed in *hh* mutant adult brains (h). Expression of *hsp40* (j) or *hsp68* (k) in the glia of *hh* mutants reduces the occurrence of aggregates. Quantification of Ubq signal intensity (l-l’) were acquired from the equivalent boxed areas (i) in all samples, box 1: CB/OL boundary and box 2: midline of central brain (h-k, S5q-r), scale bars 50μm. Area quantified for Ubq signal intensity: 31.62μm x 31.62μm (1000μm^2^). Two-way t-test used to analyse differences in Ubq signal intensity between *hh* mutant and control samples in l-l’, p<0.001***). Images (h-k) consists of a stack of 25μM (1 μM per section). (m-n) Immunoblots showing pEIF2a levels in *hh* mutant and *hh* mutant flies expressing *hsp68* in the glia in day 12 adult head samples. (n) *hh* mutant flies exhibit a 2-fold increase in pEIF2a levels, which is rescued to control levels in *hh* mutant samples with glial expression of *hsp68*, (p<0.0001****, one-way ANOVA with post-hoc Tukey).(o-p) Overexpression of *hsp68* in the glia strongly rescues the *hh* mutant short-lived phenotype, log rank test, p<0.0001 (o) and (p) *hh*-defective locomotion – two-way ANOVA test, p<0.001)

Because of the well-accepted role of Hsps in regulating proteostasis (Lindquist and Craig, 1988) we hypothesised that the decrease in *hsp40* and *hsp68* expression level that we observed in *hh* mutant adult flies may promote accumulation of protein aggregates. We tested this hypothesis using a specific antibody detecting ubiquitin-positive aggregates in the *Drosophila* brain (Nezis et al., 2008). Accordingly, we observed that old *hh* mutant flies exhibited a strikingly elevated level of ubiquitinated aggregates when compared to age-matched control flies (Fig. 4e-g, 4l-l’). Moreover, this appeared predominantly localised in midbrain glia tracts, which are enriched for the proteoglycan Perlecan (Pcan; Davie et al., 2018), and also express Ci (Fig. 4g, S5m-p). In addition, ubiquitinated aggregates were also detected in glial cells of the anterior and posterior brain regions of the *hh* mutant fly brain (Fig. S5e-h, i-l). These observations strongly suggest that a reduction in *hsp23*, *hsp40* or *hsp68* in *hh* mutant may cause the formation of ubiquitin positive protein aggregates glial cells. Accordingly, when either *hsp40* or *hsp68* were overexpressed in glial cells of aged adult *hh* mutants, they strongly reduced ubiquitinated aggregates in glial cells (Fig. 4h-l’).

An additional readout for proteostasis function is the measure of pEIF2α levels which is indicative of the presence of misfolded proteins. The <u>u</u>nfolded <u>p</u>rotein <u>r</u>esponse (UPR) in the <u>E</u>ndoplasmic <u>r</u>eticulum (ER) activates a signalling cascade which elevates pEIF2α levels (Halliday et al., 2017). We tested this using the pEIF2α antibody, which has been used to quantify pEIF2α in *Drosophila* (Edenharter et al, 2018). Interestingly we found that *hh* mutant adult brains exhibited a two-fold increase in pEIF2α levels, which are strongly reduced to wild type control levels in *hh* mutant adult flies expressing *hsp68* in the glia (Figure 4m-n). Conversely Hh overexpressing flies exhibit a 32% reduction in pEIF2α levels compared to age-matched control flies (Fig. S5t-u).

Finally, to ascertain whether *hsp40* or *hsp68* contribute to *hh*-dependent survival and fitness, we specifically expressed each of them in glial cells of *hh* mutant flies (Fig. 4o-p and Fig. S4b-d). Remarkably, we found that expression of either *hsp68* or *hsp40* potently increases the reduced lifespan observed in *hh* mutant animals (Fig 4o, S4b-c”). In contrast, *hsp23* alone was ineffective (Fig. S4d-d”). We also observed that glial expression of *hsp68* rescued the defective climbing ability of *hh* mutants by 57% (Fig. 4p). Overall, our data demonstrates that the Hh signalling in glial cells promotes expression of protective chaperones required for maintaining correct proteostasis during adult life.

### Hh signalling in the glia has a protective function in a human Amyloid beta disease model

Defects in glial proteostasis observed upon Hh depletion and their rescue via expression of Hh target chaperones in the glia, raises the possibility that stimulating Hh signalling in the glia may reduce aggregate formation and its concommitant toxicity. In *Drosophila*, expression of the human Amyloid-β 1−42 (Aβ 1−42) in the glia induces abnormal aggregate formation, reduced lifespan and defective locomotor activity (Jonson et al, 2018). Consistent with this study we expressed, in the adult *Drosophila* glia, the human arctic variant of Aβ 1−42 (Aβ 1−42 ^E693G^) which is known to have a high propensity to aggregate and form fibrils *in vitro* (Nilsberth et al, 2001). We observed the formation of widespread amyloid aggregates throughout the adult brain (Fig. S6a-h.). This phenomenon is also noticeable in the giant glial cells of the optic chiasm (dashed box – Fig. S6e and g) in the midbrain sections. The DAPI positive cells exhibit an abnormal morphology with a markedly reduced cell number in this region (Fig. S6g”) compared to age-matched wild type control, which exhibit no detectable amyloid aggregates (Fig. S6f, h-h’’). Expression of human Aβ 1−42 ^E693G^ in the glia also leads to reduction in lifespan as well as an age-dependent decline in locomotor activity (Fig. 5a-c).

**Figure 5.**
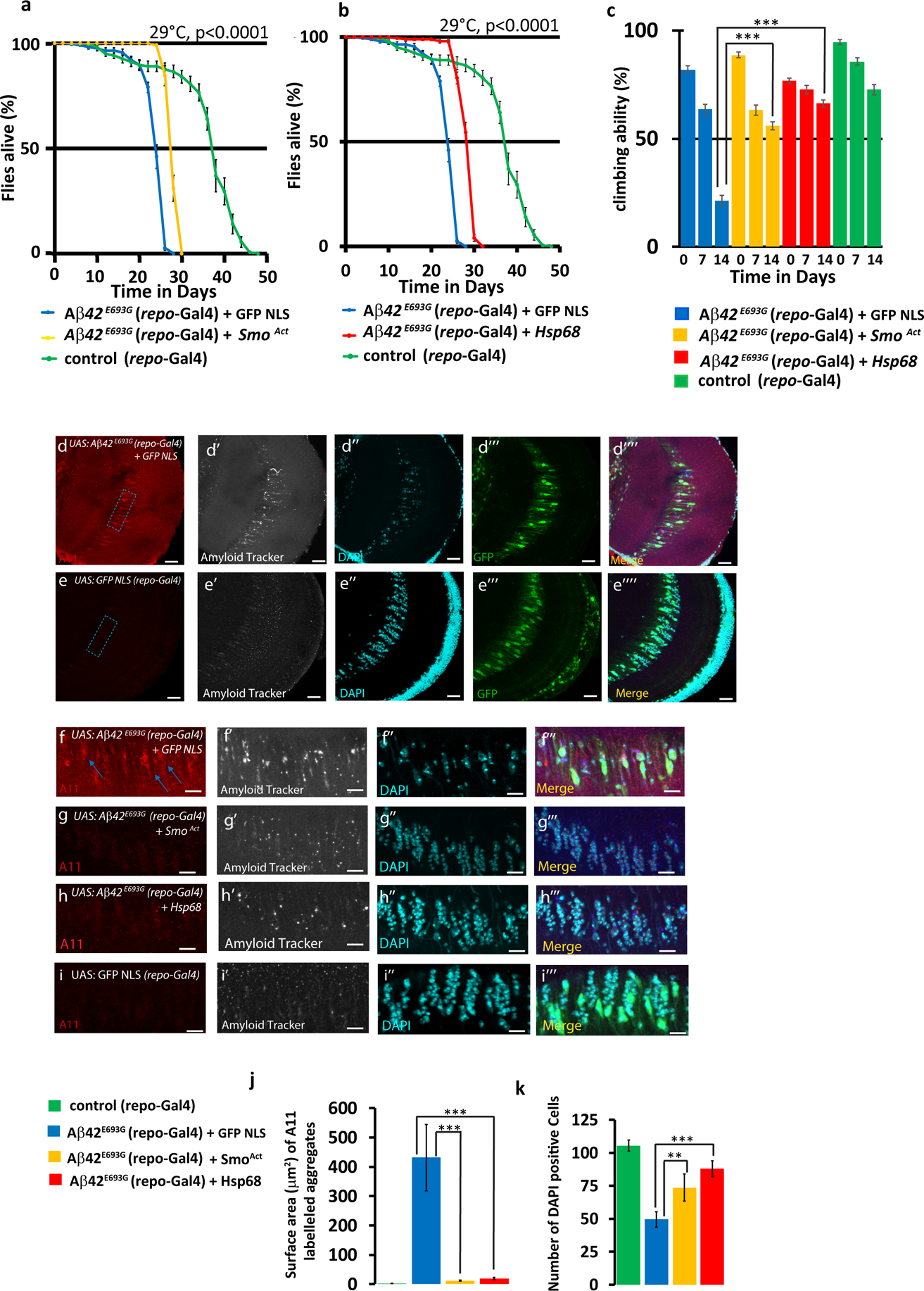
Hh signalling in the glia has a protective function in a human Amyloid beta disease model. (a-c) The shortened lifespan and debilitated locomotion that results from glial expression of human Aβ42 ^E693G^ can be partially rescued by expression of (a, c) an activated form of Smo (*Smo ^Act^*) or (b, c) *hsp68* expression in the glia. The rescues are more potent in the human Aβ42 ^E693G^ middle aged animals at the 20% mortality rate (Table S2 and S3; as for locomotion assays in c, lifespan assays in a and b were performed in the same experiment and yellow and red curves separated for visualisation). For lifespan analysis log rank test utilised (p<0.0001), for climbing activity two-way Anova test was used, p<0.001***. (d-e) Confocal section (5μm stack) of the optical lobe of 15-day old flies expressing human Aβ42 ^E693G^ in the glia, (d) showing the presence of amyloid aggregates (demarcated by the dashed box), in the giant glial cells of the optic chiasm (glial cells labelled with GFP NLS in the merged image); in control specimens the amyloid aggregates are not discernable (e), Scale bars 20μm. In order to quantify the presence of aggregates and to count cell number in the region of interest - ROI (dashed box), an area of 43.14μm x 91.96μm was selected for the ROI (f). Blue arrows indicate the presence of A11 positively stained amyloid aggregates in flies overexpressing human Aβ42 ^E693G^ in the glia– red channel (f). The grey channel displays the presence of the amyloid (f’), as detected by the Amyloid Probe (AmyTracker^TM^680) and the cyan channel displays DAPI positive cells (f”), with the merge of all three channels also shown (f’’’), Scale Bars 10 μm. Overexpression of either (g) *Smo^Ac^*^t^ (h) or *hsp68* is able to reduce the presence of amyloid aggregates (j), (see Experimental procedures for amyloid aggregate quantification), and rescue the decreased cell number (g”, h”) observed in flies overexpressing human Aβ42 ^E693G^ (k). For both amyloid aggregate quantifications and cell number the two way t-test was used (p<0.001***)

Stimulation of the Hh pathway in glial cells expressing human Aβ 1−42 ^E693G^ via expression of Smo^act^ reduced the presence of amyloid aggregates in the giant glial cells of the optic chiasm and rescued DAPI positive cells in this region (Fig. 5g-g”’, S6i-i’’’, Fig 5j-k), and significantly rescued shortened lifespan and locomotion defects (Fig. 5a, c). The rescues were more effective in the Aβ 1−42 ^E693G^ in young animals (80% survival rate) expressing Smo^Act^ with a rescue of 40% and a 69% rescue of debilitated locomotor activity at day 14 (Fig. 5a, c, Table S2-S3). Similarly, the expression of Hsp68 in the glia of Aβ 1−42 ^E693G^ expressing animals improved longevity and locomotion by 43% and 88%, respectively (Fig 5b-c, Table S2-S3) as well as a reduction in the presence of amyloid aggregates in the glia and a more potent rescue of cell number (Fig. 5h-h’’’,5j-k and S6j-j’’’).

Taken together our study suggests that activating Hh signalling in the glia is protective against abnormal proteostasis. Our proof of concept study utilizing glial expression of the human Aβ 1−42 ^E693G^ artic variant, provides an indicator for the protective capacity of Hh signalling in a disease paradigm.

## DISCUSSION

Our findings provide evidence for a previously unidentified function for the Hh morphogen during adulthood distinct from its role as a patterning organizer and a tumorigenic factor (Briscoe and Thérond, 2013). Our results suggest that Hh signalling in the glia of the adult brain controls the expression of chaperone molecules, including Hsp40 and Hsp68, which repress the formation of age-associated protein aggregates. We propose a model in which Hh morphogen signaling, interpreted by glial cells, regulates lifespan determination and associated fitness, whilst maintaining adult brain homeostasis, including DA neuron protection (Figure 6).

**Figure 6.**
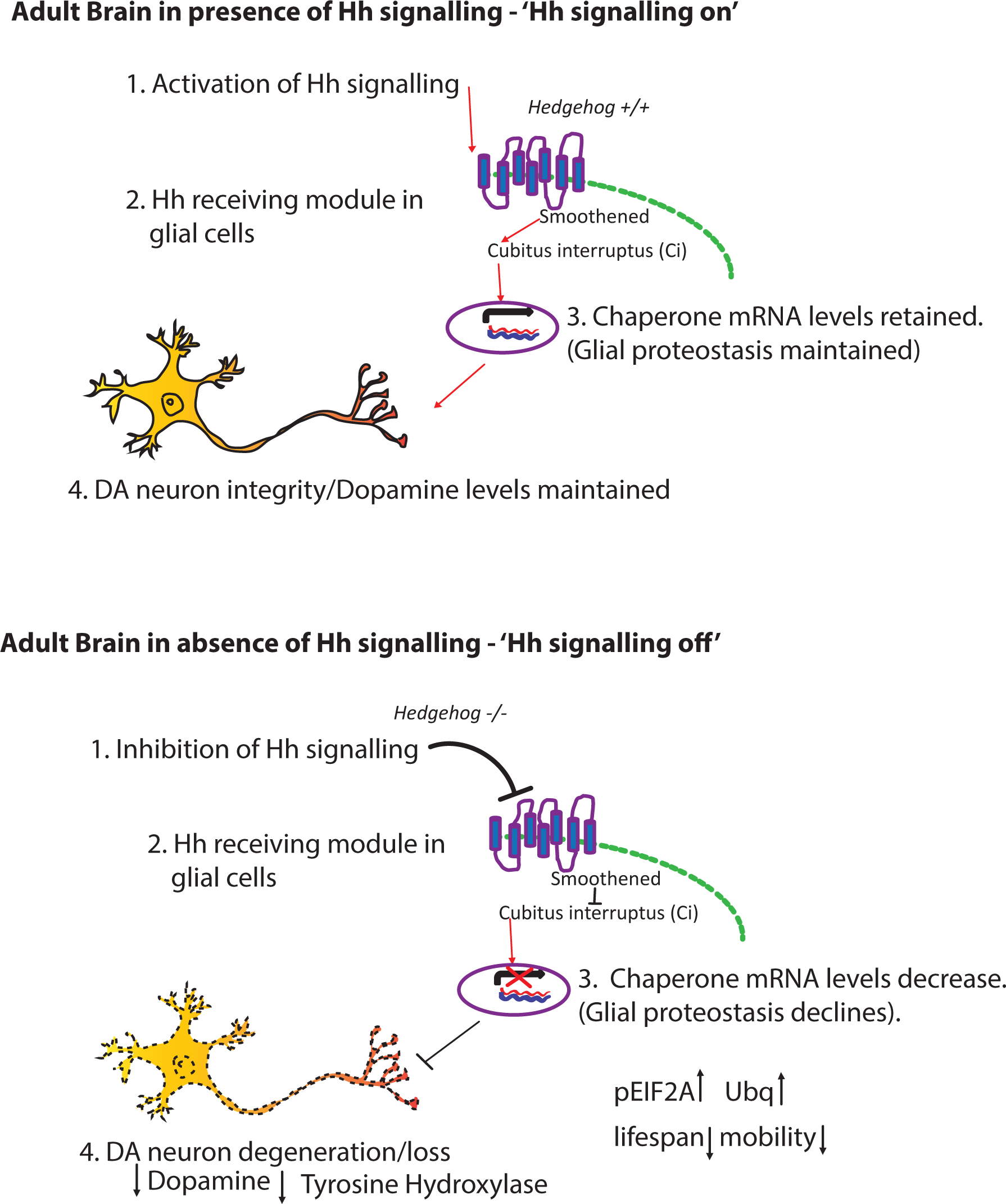
Proposed model by which Hh signalling mediates glial proteostasis and neuroprotection. We put forward the following model: a) **1.** the presence of the Hh signal **2**. induces activation of the canonical Hh signalling pathway in the glia via Ci activation. **3**. This regulates mRNA levels of downstream target chaperones in the glia, which maintain proteostasis and **4**. maintains DA neuron integrity. b) **1.** Conversely in the absence of Hh signalling, **2** Ci remains inactive in glia and **3** mRNA levels of downstream chaperones are decreased. Subsequently, glial proteostasis declines, leading to an increase in abnormal protein aggregation, characterised by an elevated level of Ubq positive aggregates. This proteotoxic environment in the glia is deleterious to lifespan and mobility of the organism and leads to **4** degeneration of DA neurons and a decrease in dopamine levels, potentially due to a reduction in neuroprotective factors produced in the glia.

### Hh signaling in glial cells as a lifespan determinant

There is an emerging body of evidence that diverse glial cell dysfunction has an impact on neuroprotection and organismal lifespan. For instance, disturbance of mitochondrial function by frataxin silencing (Navarro et al., 2010) or impairment of glycolysis via depletion of glycolytic enzymes in the glial cells reduces lifespan and induces neurodegeneration in *Drosophila* (Volkenhoff et al., 2015; Miller et al., 2016). Additionally activation of the innate immunity response, namely the IMD/NF-κΒ pathway, in glial cells resulted in an increased expression of antimicrobial peptides, inducing neurodegeneration and diminishing lifespan (Kounatidis et al., 2017). Conversely, glial specific suppression of the IMD/NF-κΒ signalling pathway culminated in an increased presence of the AKH hormone and lifespan extension (Kounatidis et al., 2017). Furthermore, it has been proposed that the glial neuropeptide RGBA-1 is able to modulate the rate of ageing in *C. elegans* (Yin et al., 2017). Overall our study reinforces the above notion that glia have an important function in the modulation of lifespan and neuroprotection.

Indeed, our study revealed that the Hh signalling pathway is a novel regulator of glial proteostasis. The Hh targets in this cell type, namely Hsp40 and Hsp68, are highly conserved chaperones, (for which the Hsp40 and Hsp70 chaperone family members are the conserved homologues in mammals). These chaperones form part of a major protein complex, which recognises and binds ubiquitinated misfolded proteins, preventing their aggregation and facilitating their refolding or degradation (Lindquist et al., 1988; Jackrel and Shorter, 2017; Zarouchlioti et al., 2018). Interestingly, Hsp40 and Hsp70 act cooperatively as safeguards to prevent both the misfolding of newly synthesized proteins and protein aggregation upon cellular stress. A third Hh target, Hsp23, classified as a small heat shock protein “holdase”, binds protein folding intermediates to prevent aggregation, but, in contrast to the Hsp40/70 complex, is unable to refold protein into their native states (Morrow et al., 2015). This functional difference could provide an explanation for the lack of rescue when Hsp23 is expressed in glia of *hh* mutant animals. Our study demonstrates that glial overexpression of Hsp40 or Hsp68 is able to rescue proteostasis defects present in the *hh* mutant glial cells, characterized by an elevated level of ubiquitin. Interestingly, *hh* mutant flies also exhibit increased levels of pEIF2α, indicative of activation of the unfolded proteins response (Halliday et al., 2017), which can be restored to wild type levels through glial expression of Hsp68. Conversely, Hh overexpressing flies display reduced levels of pEIF2α, suggesting that stimulating Hh signalling is protective against protein misfolding.

Some unanswered questions remain, for instance what subtype of glial cells are responsible for mediating Hh coordinated adult survival and neuronal integrity. One possibility is, that the cortex glia which envelopes neuronal cell bodies could perform this function. It has been suggested that this subtype can provide trophic support to neurons (Stork et al, 2012). Alternatively, the ensheathing glia or the surface glia forming the glia of the blood brain barrier may be the location of Hh mediated neuroprotection. Further investigation is required to address the contribution of each glial subtype to Hh mediated survival and fitness.

A further line of inquiry to address would be the degree to which protective Hh signalling in the glia is acting in a cell/non-cell autonomous manner. Given that both, Hsp40 and Hs68, were demonstrated in our study to reduce the presence of ubiquitin positive aggregates in the glia as well as rescue survival rates of *hh* mutant flies suggests that they act in an autonomous manner, maintaining glial proteostasis. A recent study however demonstrated that both, Hsp40 and Hsp70, can be secreted on exosomes and exert a paracrine neuroprotective effect in an *in vitro* co-culture experiment (Takeuchi et al., 2015). This suggests that the chaperone targets of Hh signalling may be secreted from the glia and thereby contribute towards Hh-mediated neuroprotection and lifespan determination in a non-cell autonomous manner.

### The potential of Hh signaling in alleviating degenerative disease

As proof of principle we tested components and targets of Hh signalling in an *in vivo* Alzheimer’s disease (AD) model in *Drosophila* expressing the human arctic variant of Aβ 1−42 (Aβ 1−42 ^E693G^) in the glia. In agreement with a recent study (Jonson et al., 2018), here we show that glial expression of the Aβ 1−42 ^E693G^ variant shortened lifespan, induced debilitated locomotion and increased the presence of amyloid aggregates. Remarkably, we found that induction of Hh signalling in the glia was able to considerably rescue the Aβ 1−42 ^E693G^ – dependent phenotype, both at the molecular and physiological level, providing the first evidence to our knowledge to confirm the efficacy of Hh signalling in an *in vivo* AD model. We also show that increasing expression of Hsp68 reduces the level of Aβ 1−42 ^E693G^ aggregates *in vivo*, which is consistent with a study showing that expression of Hsp70 reduces the level of amyloid β aggregates in transgenic mice brains expressing a variant of the amyloid precursor protein (Hoshino et al., 2011). Overall we propose that targets under the regulation of Hh signaling in the glia have the potential to ameliorate proteinopathies, such as those caused by Aβ 1−42 in Alzheimer’s disease. It is now proposed that the glia play an important role in various proteinopathies and could function as carriers for the initiation and propagation of aggregates in the adult brain. In fact it has been demonstrated that glial cells can replicate and seed Aβ amyloid fibrils in the mammalian brain independent of neuronal expression (Veeraraghavalu et al., 2014). Thus, activation of the Hh signalling pathway in the glia may have beneficial effects in combating a wide-range of proteinopathies which manifest themselves through aberrant protein aggregation and misfolding as observed in Parkinson’s, Alzheimer and Huntington diseases.

### A conserved role for Hh signaling in healthspan and neuroprotection

We propose that this new function for Hh signaling as a lifespan determinant may be conserved in mammals. All molecular players described here are conserved; the vertebrate counterpart of *Drosophila* Hh, Shh, is also expressed in mouse adult neurons and glial cells respond to this signal (Garcia et al., 2010, Farmer et al., 2016). Since the activation of Shh signalling protects against diverse neurotoxins such as amyloid beta peptide *in vitro* (Yao et al., 2017), our study suggests that Hh signaling in glia may ameliorate age-related neurodegenerative diseases caused by aberrant protein aggregation as well as prolong healthspan and fitness in mammals.

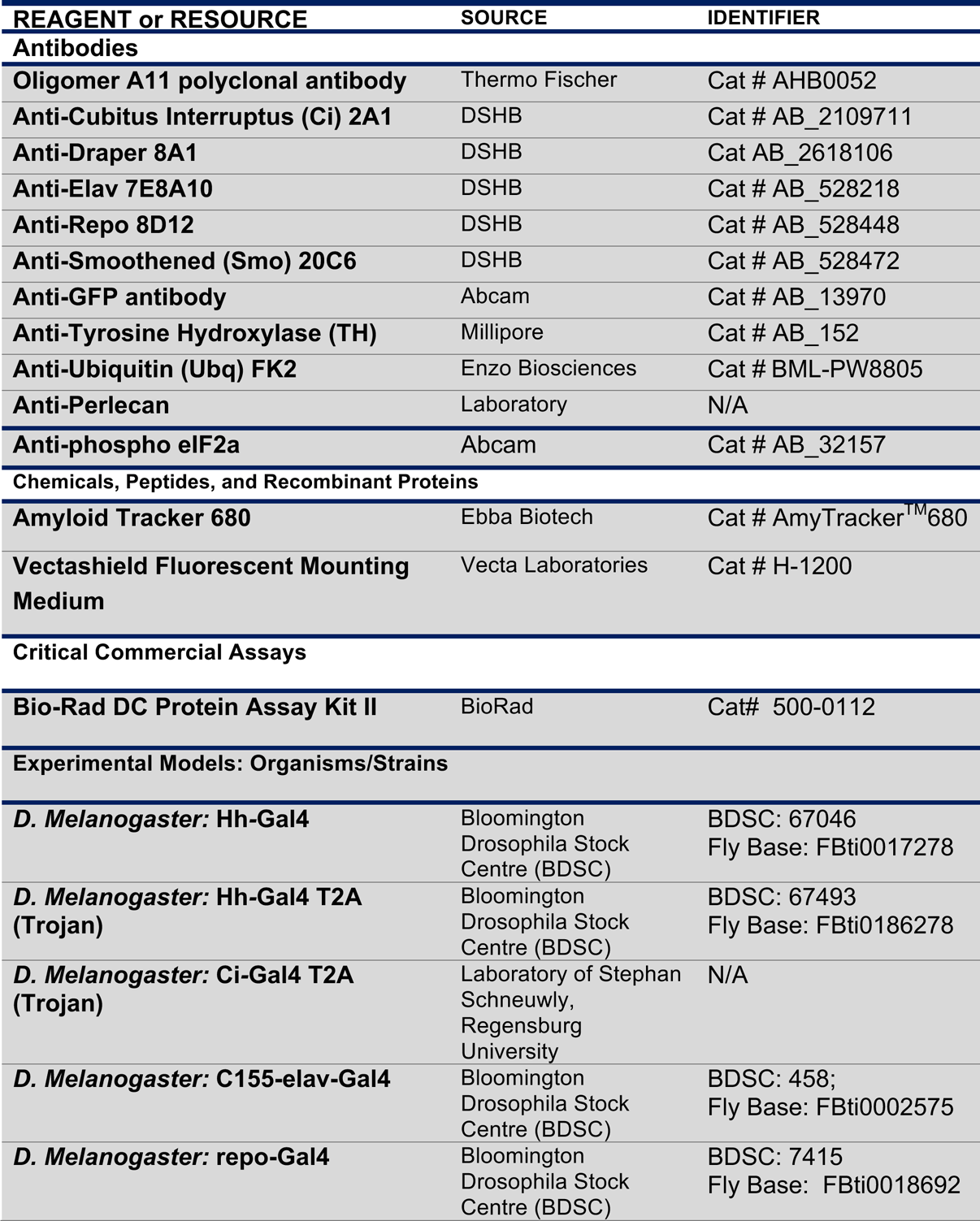
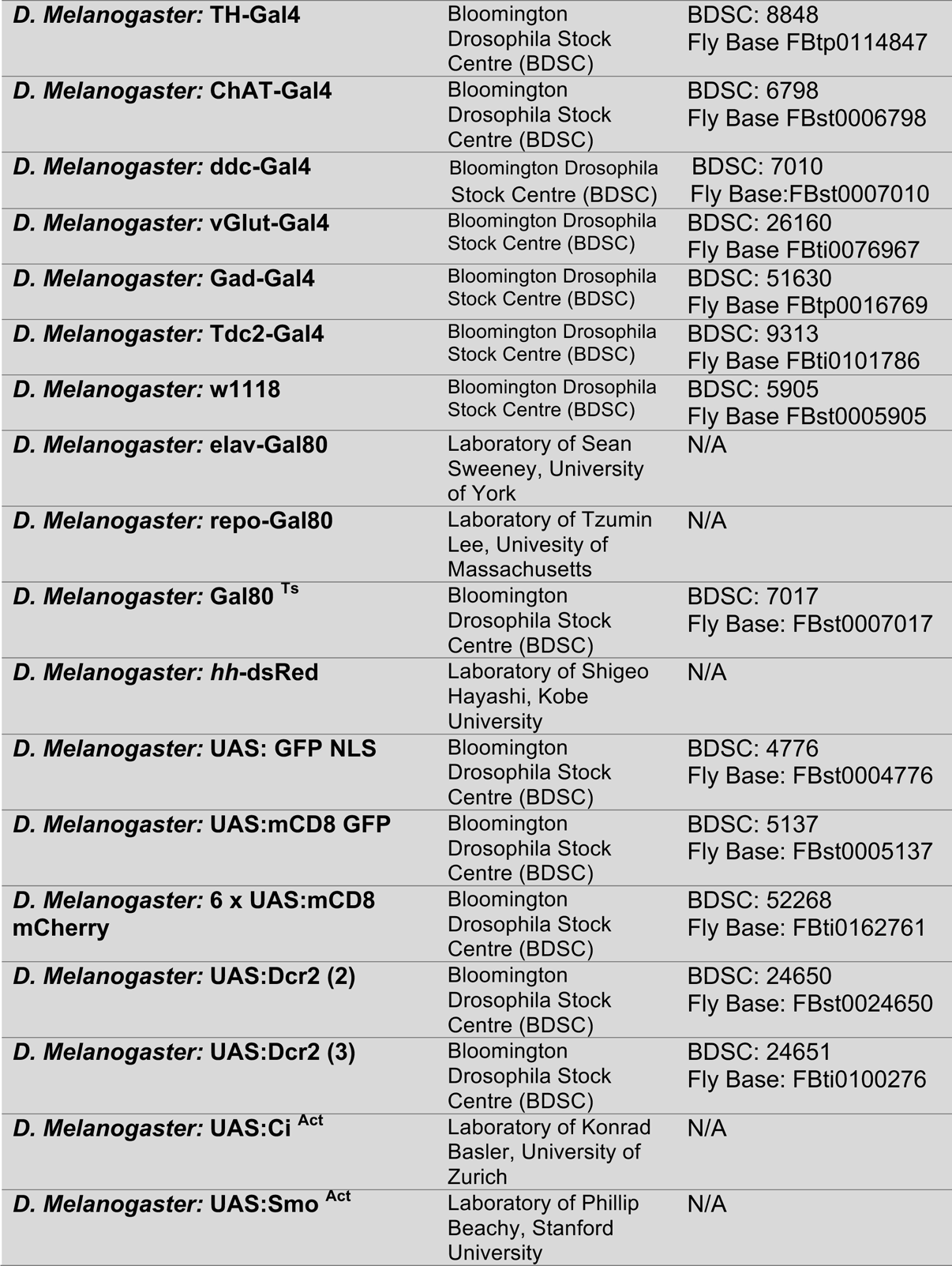
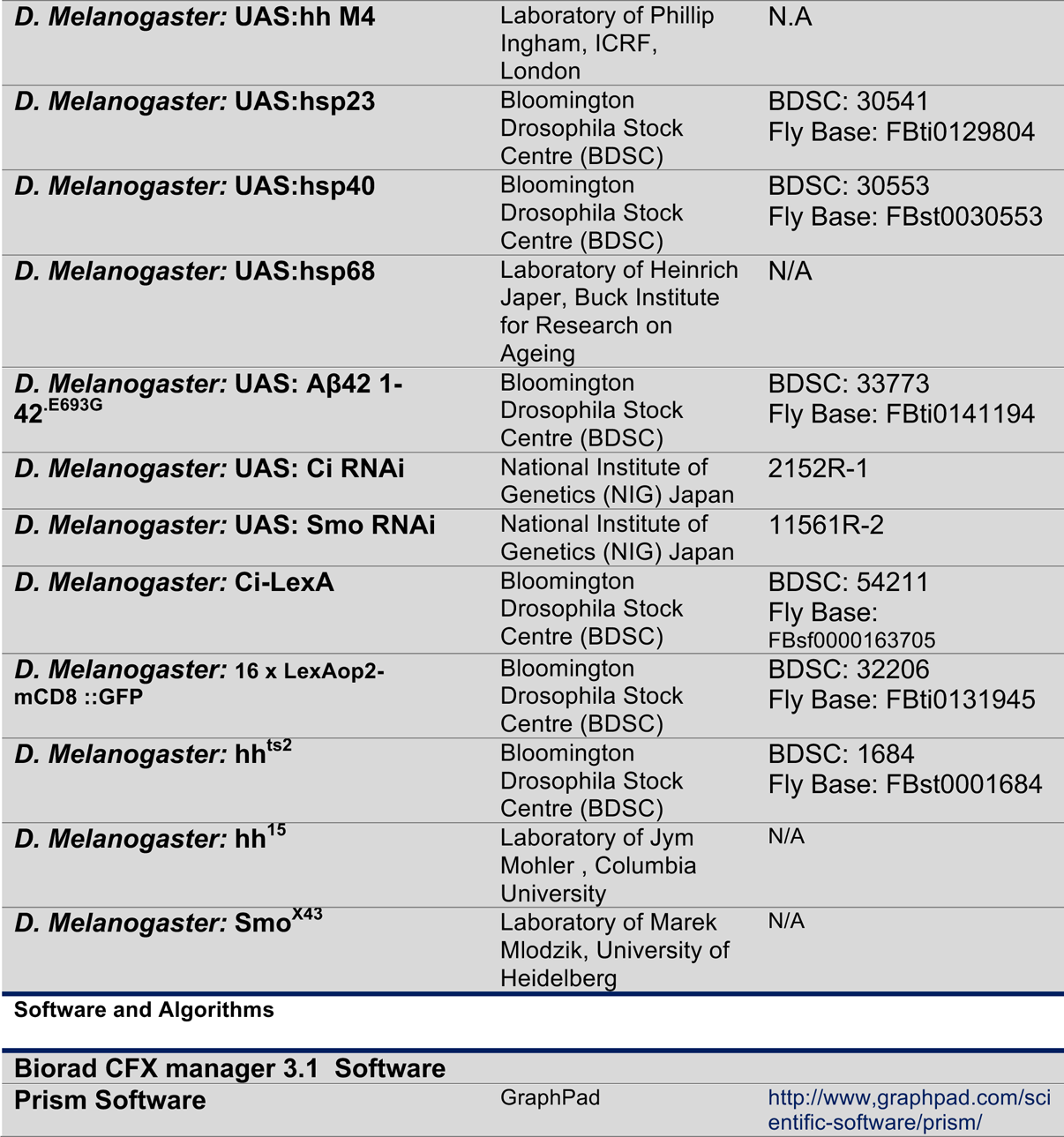
**KEY RESOURCES TABLE**

KEY RESOURCES TABLE
| REAGENT or RESOURCE | SOURCE | IDENTIFIER |
| --- | --- | --- |
| **Antibodies** |  |  |
| **Oligomer A11 polyclonal antibody** | Thermo Fischer | Cat # AHB0052 |
| **Anti-Cubitus Interruptus (Ci) 2A1** | DSHB | Cat # AB_2109711 |
| **Anti-Draper 8A1** | DSHB | Cat AB_2618106 |
| **Anti-Elav 7E8A10** | DSHB | Cat # AB_528218 |
| **Anti-Repo 8D12** | DSHB | Cat # AB_528448 |
| **Anti-Smoothed (Smo) 20C6** | DSHB | Cat # AB_528472 |
| **Anti-GFP antibody** | Abcam | Cat # AB_13970 |
| **Anti-Tyrosine Hydroxylase (TH)** | Millipore | Cat # AB_152 |
| **Anti-Ubiquitin (Ubq) FK2** | Enzo Biosciences | Cat # BML-PW8805 |
| **Anti-Perlecan** | Laboratory | N/A |
| **Anti-phospho eIF2a** | Abcam | Cat # AB_32157 |
| **Chemicals, Peptides, and Recombinant Proteins** |  |  |
| **Amyloid Tracker 680** | Ebba Biotech | Cat # AmyTracker™680 |
| **Vectashield Fluorescent Mounting Medium** | Vecta Laboratories | Cat # H-1200 |
| **Critical Commercial Assays** |  |  |
| **Bio-Rad DC Protein Assay Kit II** | BioRad | Cat# 500-0112 |
| **Experimental Models: Organisms/Strains** |  |  |
| ***D. Melanogaster*: Hh-Gal4** | Bloomington Drosophila Stock Centre (BDSC) | BDSC: 67046<br>Fly Base: FBti0017278 |
| ***D. Melanogaster*: Hh-Gal4 T2A (Trojan)** | Bloomington Drosophila Stock Centre (BDSC) | BDSC: 67493<br>Fly Base: FBti0186278 |
| ***D. Melanogaster*: Ci-Gal4 T2A (Trojan)** | Laboratory of Stephan Schneuwly, Regensburg University | N/A |
| ***D. Melanogaster*: C155-elav-Gal4** | Bloomington Drosophila Stock Centre (BDSC) | BDSC: 458;<br>Fly Base: FBti0002575 |
| ***D. Melanogaster*: repo-Gal4** | Bloomington Drosophila Stock Centre (BDSC) | BDSC: 7415<br>Fly Base: FBti0018692 |
| ***D. Melanogaster*: TH-Gal4** | Bloomington Drosophila Stock Centre (BDSC) | BDSC: 8848<br>Fly Base FBtp0114847 |
| ***D. Melanogaster*: ChAT-Gal4** | Bloomington Drosophila Stock Centre (BDSC) | BDSC: 6798<br>Fly Base FBst0006798 |
| ***D. Melanogaster*: ddc-Gal4** | Bloomington Drosophila Stock Centre (BDSC) | BDSC: 7010<br>Fly Base:FBst0007010 |
| ***D. Melanogaster*: vGlut-Gal4** | Bloomington Drosophila Stock Centre (BDSC) | BDSC: 26160<br>Fly Base FBti0076967 |
| ***D. Melanogaster*: Gad-Gal4** | Bloomington Drosophila Stock Centre (BDSC) | BDSC: 51630<br>Fly Base FBtp0016769 |
| ***D. Melanogaster*: Tdc2-Gal4** | Bloomington Drosophila Stock Centre (BDSC) | BDSC: 9313<br>Fly Base FBti0101786 |
| ***D. Melanogaster*: w1118** | Bloomington Drosophila Stock Centre (BDSC) | BDSC: 5905<br>Fly Base FBst0005905 |
| ***D. Melanogaster*: elav-Gal80** | Laboratory of Sean Sweeney, University of York | N/A |
| ***D. Melanogaster*: repo-Gal80** | Laboratory of Tzumin Lee, University of Massachusetts | N/A |
| ***D. Melanogaster*: Gal80<sup>Ts</sup>** | Bloomington Drosophila Stock Centre (BDSC) | BDSC: 7017<br>Fly Base: FBst0007017 |
| ***D. Melanogaster*: hh-dsRed** | Laboratory of Shigeo Hayashi, Kobe University | N/A |
| ***D. Melanogaster*: UAS: GFP NLS** | Bloomington Drosophila Stock Centre (BDSC) | BDSC: 4776<br>Fly Base: FBst0004776 |
| ***D. Melanogaster*: UAS:mCD8 GFP** | Bloomington Drosophila Stock Centre (BDSC) | BDSC: 5137<br>Fly Base: FBst0005137 |
| ***D. Melanogaster*: 6 x UAS:mCD8 mCherry** | Bloomington Drosophila Stock Centre (BDSC) | BDSC: 52268<br>Fly Base: FBti0162761 |
| ***D. Melanogaster*: UAS:Dcr2 (2)** | Bloomington Drosophila Stock Centre (BDSC) | BDSC: 24650<br>Fly Base: FBst0024650 |
| ***D. Melanogaster*: UAS:Dcr2 (3)** | Bloomington Drosophila Stock Centre (BDSC) | BDSC: 24651<br>Fly Base: FBti0100276 |
| ***D. Melanogaster*: UAS: Ci<sup>Act</sup>** | Laboratory of Konrad Basler, University of Zurich | N/A |
| ***D. Melanogaster*: UAS:Smo<sup>Act</sup>** | Laboratory of Phillip Beachy, Stanford University | N/A |
| ***D. Melanogaster*: UAS:hh M4** | Laboratory of Phillip Ingham, ICRF, London | N.A |
| ***D. Melanogaster*: UAS:hsp23** | Bloomington Drosophila Stock Centre (BDSC) | BDSC: 30541<br>Fly Base: FBti0129804 |
| ***D. Melanogaster*: UAS:hsp40** | Bloomington Drosophila Stock Centre (BDSC) | BDSC: 30553<br>Fly Base: FBst0030553 |
| ***D. Melanogaster*: UAS:hsp68** | Laboratory of Heinrich Japer, Buck Institute for Research on Ageing | N/A |
| ***D. Melanogaster*: UAS: Aβ42 1-42<sup>E693G</sup>** | Bloomington Drosophila Stock Centre (BDSC) | BDSC: 33773<br>Fly Base: FBti0141194 |
| ***D. Melanogaster*: UAS: Ci RNAi** | National Institute of Genetics (NIG) Japan | 2152R-1 |
| ***D. Melanogaster*: UAS: Smo RNAi** | National Institute of Genetics (NIG) Japan | 11561R-2 |
| ***D. Melanogaster*: Ci-LexA** | Bloomington Drosophila Stock Centre (BDSC) | BDSC: 54211<br>Fly Base: FBsf0000163705 |
| ***D. Melanogaster*: 16 x LexAop2-mCD8 ::GFP** | Bloomington Drosophila Stock Centre (BDSC) | BDSC: 32206<br>Fly Base: FBti0131945 |
| ***D. Melanogaster*: hh<sup>ts2</sup>** | Bloomington Drosophila Stock Centre (BDSC) | BDSC: 1684<br>Fly Base: FBst0001684 |
| ***D. Melanogaster*: hh<sup>15</sup>** | Laboratory of Jym Mohler, Columbia University | N/A |
| ***D. Melanogaster*: Smo<sup>x43</sup>** | Laboratory of Marek Mlodzik, University of Heidelberg | N/A |
| Software and Algorithms |  |  |
| **Biorad CFX manager 3.1 Software** |  |  |
| **Prism Software** | GraphPad | http://www.graphpad.com/scientific-software/prism/ |

### EXPERIMENTAL PROCEDURES

#### Fly Stocks and Lifespan/Locomotion Assays

Flies were raised on standard media at 18°C for all experiments and transferred to 29°C, no longer than 24 hours after adult eclosion, to ensure adult specific inactivation of the *hh* mutants and adult specific expression of the relevant transgenes of the Hh signalling pathway using the Gal80^ts^/UAS/GAL4 system during adulthood as well as to avoid developmental defects prior to adult eclosion. Stock lines and GAL4 driver lines were obtained from the *Drosophila* Stock center at Bloomington, or are described in the methods text. Flies were generated or backcrossed a minimum of 5 generations into a controlled uniform homogeneous genetic background (Flybase ID FBst0005905, *w111*8), in order to ensure that all phenotypes were robust and not associated with variation in genetic background. In this uniform homogeneous genetic background, the survival time of control flies is highly reproducible. Startled-induced negative geotaxis was used to assess fly locomotion. To determine lifespan, newly eclosed females were mated for 24 to 48 hours at 18°C collected and maintained at 20 flies per vial and transferred to fresh 11 vials every two days while being scored for survival. A minimum of 180 to 300 flies were used per genotype per lifespan assay, unless otherwise indicated.

#### Immunohistochemistry

Adult male heads were dissected and immunostained with the specified antibodies using standard Procedures (Wu and Luo, 2006).

## Statistical Analysis

Survival assays were analyzed in Excel (Microsoft) and in Prism software (GraphPad) for survival curves and statistics.

### qRT-PCR analysis and Western Blots

Techniques of molecular biology, and histology were standard. Expression of genes from fly heads extracts was carried out using standard procedures for mRNA extraction, cDNA synthesis and semi-quantitative Real Time PCR.

## Supplementary Information

Supplemental information includes Supplemental Experimental Procedures, six figures and six tables can be found with this article:

### Supplemental Experimental Procedures Genetic background

All flies were generated in the same uniform homogeneous genetic background in order to prevent genetic background effects, (line 5905 (Flybase ID FBst0005905, *w111*8)), and backcrossed a minimum of 5 generations into this uniform *w1118* genetic background. This ensures that the observed phenotypes were due to gene manipulations, and not variation in genetic background. In these carefully controlled experiments, the lifespan of control flies was highly uniform and repeatable when 180 or more individuals were used for lifespan analysis (see Table S1). The short hand and the corresponding full-genotypes as well as explanation of the components utilised for the lifespan/mobility experiments are listed in Table S4.

### Fly stocks and genetics

The UAS/GAL4/GAL80^ts^ system was utilised (Brand and Perrimon., 1993) to drive adult tissue-specific protein expression of the relevant UAS: transgenic lines and to avoid developmental defects prior to adult eclosion. Gal80^ts^ is an effective repressor of Gal4 activity at 18°C. At 25°C or 29°C Gal80^ts^ is partially or completely inhibited, allowing Gal4 activity thereby driving the expression of the relevant UAS transgene. The following Gal4/Gal80 strains were used in this study: *hh-Gal4, elav-Gal4, repo-Gal4, TH-GAL4, ChAT-GAL4, Ddc-Gal4, vGlut-GAL4, Gad-GAL4, Tdc2-GAL4, w1118, elav-Gal80, Gal80^ts^* (Bloomington, Drosophila Stock Centre) and *repo-Gal80* (a gift from T. Lee, Awakaki and Lee, 2011) *hh*-dsRed (a gift from S. Hayashi, Akimoto et al, 2005). The following UAS transgenic lines were used: *UAS:GFP NLS, UAS:mCD8::GFP, UAS: hh M4, UAS: ci^act^, UAS: smo Glu ^(Act)^* (Zhang et al, 2004)*, UAS: Dcr2, UAS: hsp23, UAS: hsp40* (Bloomington Drosophila Stock Centre) and *UAS: Hsp68* (a gift from H. Jasper; Wang et al, 2005) The *hh* mutant allele strains used were as follows: *hh^ts2^,* - a temperature sensitive allele shown to be null when inactivated at the non-permissive temperature 29°C, (Stringini et al, 1997; Palm et al, 2013; Ranieri et al, 2012) and *hh^15^*, is a presumed null allele (Mohler et al, 1988). The RNAi lines used in this study were: *Ci RNAi* (2125R-1), *Smo RNAi* (11561R-2), from fly stocks of national institute of genetics (NIG). The following lexA/LexOp line were used in this study *ci*-lexA, 16XLexAop2-mCD8::GFP (Bloomington Drosophila stock Centre). The *smo^IIX43^* mutant was a gift from Marek Mlodzik (Nüsslein-Volhard et al, 1994).

### Adult specific inactivation of the Hh mutants

To assess the role of Hh signalling in lifespan determination, we initially examined the temperature sensitive *hh* mutant (*hh^ts2^*) strain, which can be considered a *hh* null (Stringini et al, 1997; Palm et al, 2013; Ranieri et al, 2012) when placed at non-permissive temperature 29°C for a minimum of 24 h. We raised the *hh^ts2^* flies at a permissive temperature of 18°C to ensure that eclosed adult flies exhibited no development defects and were viable.

### Lifespan assay

Fly stocks were grown under standard conditions. For adult lifespan assays, parental fly lines were mated at 18°C for 48 hours with 20 females and 40 males in 177ml polypropylene bottles containing 50ml of standard 1x yeast food. Males were then removed and the remaining females were allowed to lay embryos within a 24-hour time frame. Female adult progeny were collected at 18°c and divided into cohorts of 20 per vial (unless otherwise specified) and mated with control *w1118* control males in a 1:3 ratio for a period of a maximum of 48 hours. Males were subsequently removed and cohorts of 20 mated females per vial were transferred to an 29°C incubator, with the mortality being measured approximately at the same time of day every 2 days.

### Adult locomotion ability

Startled-induced negative geotaxis was used to assess fly locomotion. To perform this negative geotaxis assay, groups of 10 adult female flies of indicated age were transferred into a polystyrene round-bottom vials (25mm x 95mm) and left to recover at room temperature for 5 minutes. Climbing ability was scored as the percentage of flies failing to climb higher than 5cm, unless otherwise stated (Table S3) from the bottom of the tube, within 10 seconds after the base of the tube was gently hit. Three repeats were carried out for each set of 10 flies and the results averaged. Between each experiment within the same set, flies were given 1 minute to recover prior to the next negative geotaxis experiment and subsequent measurement of climbing ability. For each genotype at a given age, a minimum of 100 flies were examined.

### Generation of ci-T2A-GAL4 line

In order to overcome the lack of reliable Gal4 lines for *ci*, we generated a new stock using the Trojan Gal4 expression module (T2A-Gal4 cassette). In this approach, the native gene product is cleaved and the translation of GAL4 occurs as an independent protein (Diao et al, 2015). The Trojan module was integrated into the third exon of the *ci* locus. Two constructs were generated to integrate the construct into the fly genome. Firstly, a construct to express the guide RNA (gRNA) for the targeted cleavage of the DNA at the insertion site, and secondly the construct containing the Trojan cassette flanked by homology arms necessary for integration through homology directed repair. Primers to generate the gRNA (CigRNApU6fw: 5’-CTTCGCATTCCATCTTGCCGGACT-3’ and CigRNApU6rv: 5’-AAACAGTCCGGCAAGATGGAATGC-3’) were annealed using T4 polynucleotide kinase and cloned into BbsI-digested pU6-BbsI-chiRNA vector (Addgene plasmid 45946) as described in www.crisprflydesign.org. The two homology arms were amplified from genomic DNA using the following primers (Hom1ciAgeIfw: 5’-GGCCACCGGTCATGACCAGTGCAACGCC-3’ and Hom1ciSacIIrv: 5’-GGCCCCGCGGCCCTCGCCGGCAAGATGGAATGCTG-3’, hom2ciKpnIfw: 5’-GCGCGGTACCACTTGGTTTGGGATCAGCAGCTGATTT-3’ and hom2ciSpeIrv: 5’-GCGCACTAGTCTCAATGCGACAGCAGCTACGCC-3’) and cloned into the empty backbone of the pTGEM(0) vector (Addgene plasmid 62891) using AgeI and SacII or KpnI and SpeI, respectively. A solution containing 100 ng/µl of the gRNA in pU6 and 500 ng/µl of pT-GEM(0) was injected into the vas-Cas9 line (number 51324 from Bloomington Stock Center). Positive transformants were identified by the presence of the 3XP3-dsRed module also contained in the pT-GEM vector. The vas-Cas9 chromosome marked with 3xP3-GFP were excluded during the generation of the final stock.

### Molecular Biology Real-Time PCR

Total RNA was extracted from 50 heads using peqGold TriFast reagent (PEQLAB Biotechnologie GMBH, Erlangen, Germany) following the manufacturer’s instructions. 500 ng of mRNA were converted to cDNA using QuantiTect Rev. Transcription Kit (Qiagen GmbH, Hilden, Germany) then used for qPCR with ORA qPCR Green ROX L Mix (HighQu, Kralchtal, Germany) on a CFX connect Real-Time PCR Detection System (Bio-rad, Hercules, California, USA). The ribosomal protein 49 (*rp49*) was used as internal control. The results from at least four independent biological replicates were analysed with the Bio-Rad CFX manager 3.1 software. Gene expression levels were referred to the internal control, the relative quantification was carried out by means of the ΔΔCt method and the results were expressed as relative mRNA expression. Each experiment consisted of 4-5 independent biological replicates. The genes and the pairs of primers used for the analysis are listed below.

### Pairs of primers used for qRT PCR analysis

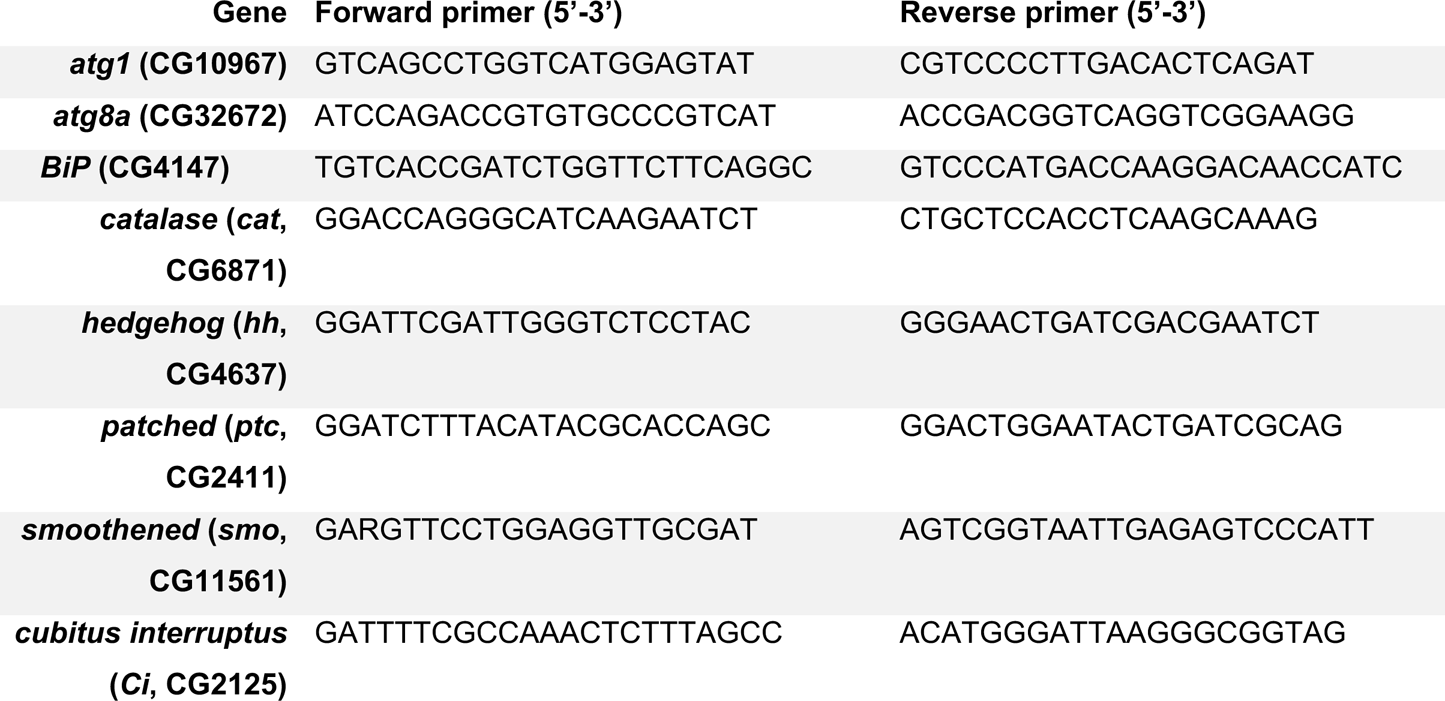
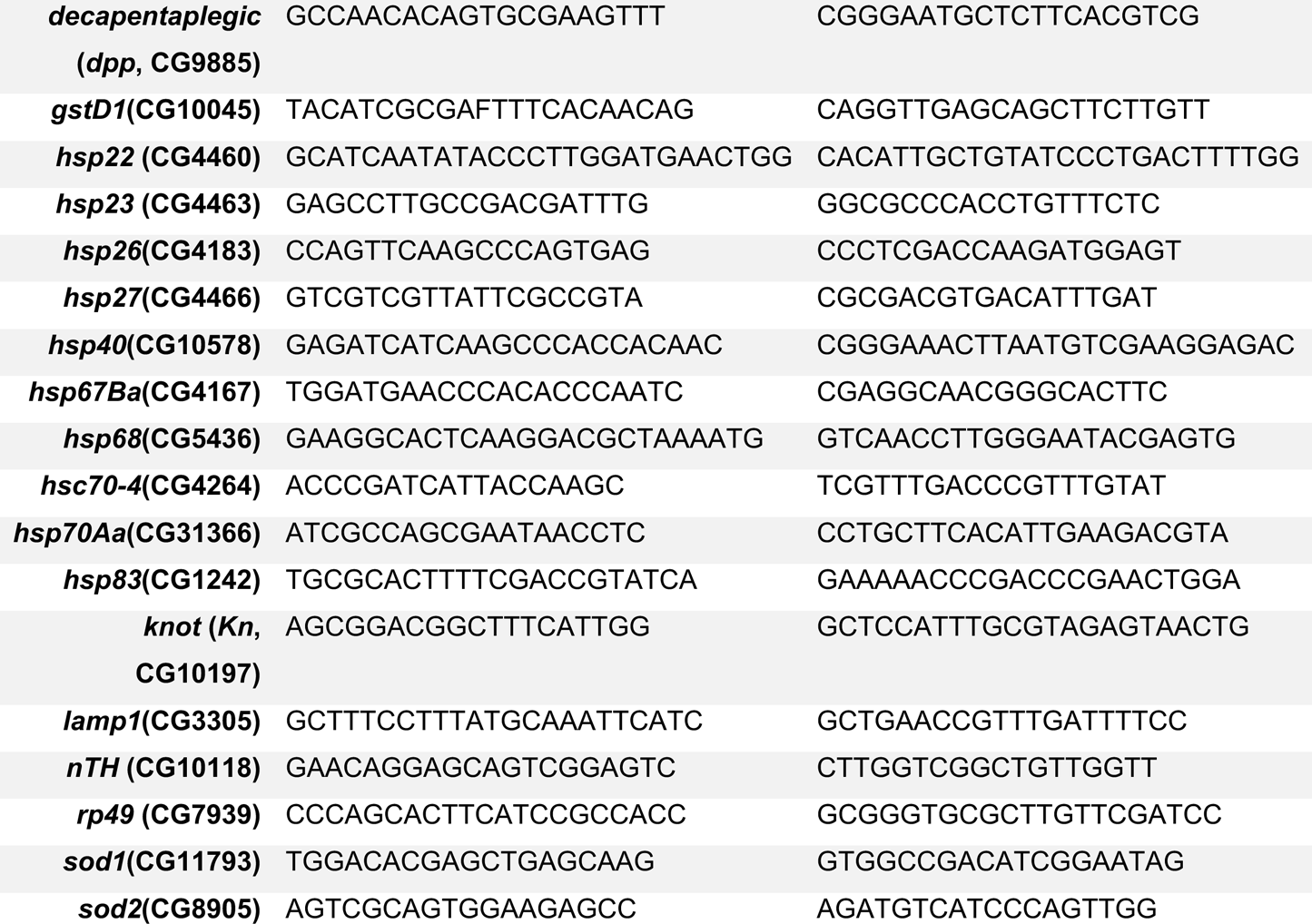

#### Immunohistochemisty

Fly brains were dissected at various time points during adulthood as specified and stained as previously described (Wu and Luo, 2006). For visualisation of dopaminergic neurons in the adult brain, the following antibody was used: rabbit anti-TH (millipore). For labelling of the neuron and glia nuclei, mouse anti-Elav: 1:50 (7E8A10 DSHB) and mouse anti-Repo: 1:50 (8D12 DSHB) were used respectively. For labelling of the glial cytosol/membrane, mouse anti-Draper: 1:50 (8A1 DSHB) was used (Purice et al, 2016). For labelling glia, rabbit anti-Perlecan 1:100 was used (gift by S.Pizette). Additional antibodies were also used: chicken anti-GFP 1:1000 (Abcam). For examining localisation of Hh signalling components in the adult brain, the following antibodies were used: mouse anti-Smo 1:50 (20C6 DSHB) and rat anti-Ci155 1:10 (2A1 gift from R. Holmgren). For labelling of aggregates in the adult brain: mouse anti-Ubq 1:500 (Enzo Biosciences) was used and for amyloid aggregates, rabbit anti-A11 (Themo Fischer) was utilised. Secondary antibodies used were as follows: goat anti-chicken Alexa488 (1:500; Invitrogen A11039); donkey anti-mouse Alexa488 (1:500; Invitrogen A21202). donkey anti-mouse Alexa546 (1:500; Invitrogen A10036). goat anti-rabbit Alexa546 (1:500; Invitrogen A11035), goat anti-rat Alexa546 (1:500; Invitrogen A11081), goat anti-mouse 647 (1:200: Invitrogen A21236) and adult brains were mounted in Vectasheld (Vecta Laboratories). Samples were embedded in Vectashield IN mounting medium (Vector Laboratories, Burlingame, CA, USA).

In each experiment, at least 4-12 flies were scanned per genotype using the Leica TSC SP5 and SP8 Confocal Laser Scanning Platforms (Leica, Germany). The detector gain and laser settings calibrated at the same level for each experiment.

#### Western Blot

For Western blots, 15 adult bodies or 40 to 50 heads were homogenized in 80 ml cold RIPA buffer (Sigma-Aldrich, St. Louis, MO, USA) with complete protease and phosphatase inhibitor mixture (Roche, Mannheim, Germany). The protein levels were determined using the BCA protein quantification kit (Thermo Scientific, Schwerte, Germany). Samples were diluted 1:4 with standard 4_Laemmli buffer, boiled for 5 min and approximately 25–30 µg of proteins were separated on 5% stacking-10% separating SDS polyacrylamide gels. The resolved proteins were electroblotted to a Protran BA 85 nitrocellulose membrane (Hartenstein, Würzburg, Germany) and were probed using rabbit anti-phospho eIF2aS1 (1:1500, ab32157, abcam, Cambridge, UK). Mouse anti-a-tubulin (1:5000, T9026, Sigma-Aldrich, St. Louis, MO, USA) was used as loading controls. Secondary fluorescent goat anti-rabbit 680RD and goat anti-mouse 800CW (1:10000, Li-Cor Inc., Bad Homburg, Germany) were used in all cases. Detection and quantification was conducted using the Odyssey system (Li-Cor Inc., Lincoln, NE, USA). Arbitrary fluorescent units were normalized to the internal loading control. Data were expressed as percentage of average control values and plotted as relative protein levels.

### DA neuron volume quantification in the adult brain with Imaris software

Dopaminergic neurons were recognised through specific labelling with either mCD8::GFP or the anti-TH immunostaining. We used the IMARIS 8.4.1 module (Bitplane Inc) to process the raw files, which consisted of the entire 3D Z stack of the adult brain. The IMARIS ‘add new surfaces’ option was selected for the relevant channel and the threshold was adjusted until all mCD8::GFP tagged neurons or neurons positive for anti-TH immunostaining were labelled as a discrete surface. Neurons were clearly distinguished from background due to the contiguous nature of their axons. We then measured the volume of DA neurons in each adult brain, using the ‘statistics’ option. All subsequent DA neuron volume data for each adult brain sample were subsequently exported to Excel. The collated data were further analysed within Prism software (GraphPad) to statistically compare *hh* mutant and age-matched control samples.

The DA neuron volume network was analysed by labelling their cell membrane with the mCD8-GFP reporter transgene using the *Ddc*-GAL4 driver in tandem with staining for TH. The *hh* mutant and control samples containing the *Ddc-*Gal4 *driver* exhibited extended survival times as a result we quantified DA neuron volume at different time points, young (0 to 7 days), middle-aged (day 21) and more aged time points (day 35). The decrease in TH immunostaining levels characteristic of older *hh* mutant flies was more delayed and manifested at latter time points. The above variation in lifespan due to the presence of the *Ddc*-Gal4 driver, is consistent with studies demonstrating that *Ddc* has been implicated as a lifespan determinant (De Luca, et al, 2003; Paaby and Schmidt; 2009)

### DA neuron cell counting

PAM DA neuron clusters were counted with the Fiji cell counting tool in a Z-projection covering the region were PAM DA neuron cell bodies were present.

### Dopamine quantifications

Dopamine levels in *Drosophila* brain extracts were determined by HPLC coupled to electrochemical detection essentially as described (Hardie and Hirsch, 2006) using a mobile phase containing 4 mM decanesulfonic acid, 50 mM citrate/acetate, pH 4.5 and 20% acetonitrile. The HPLC system consisted of a Jasco model PU-2089 Plus isocratic pump (Jasco Inc., Easton, MD, USA) coupled to an ED703 Pulse electrochemical detector (GL Sciences, Shinjuku-ku, Tokyo, Japan) equipped with a low-volume flow cell. Separation was performed with a 10 cm x 4.6 mm Hypersil H3ODS-100A column (Hichrom, Berkshire, UK) protected by a Hypersil guard cartridge. The mobile phase was delivered at 0.3 ml/min flow rate. The brains of 10 flies per genotype were dissected and directly placed into 50 μl of ice-cold 50 mM citrate/acetate, pH 4.5. They were quickly homogenized with a Teflon Pestle (Dominique Dutscher, Brumath, France). The homogenates were run through a nylon 0.2 μm centrifugal filter (VWR, Radnor, PA, USA) before analysis. Dopamine detection was performed at a detector potential of +1,000 mv relative to a conductive diamond electrode. The injection volume was 10 μl. Three independent determinations were done for each genotype and three injections were done for each determination. Results are presented as mean ± SEM.

### Ubiquitin signal intensity quantifications

Quantifications of Ubq signal intensity were acquired utilising Image J>Measurement function, from two different regions of the adult fly brain, labelled 1 CB (Central Brain)/OL (Optical Lobe) boundary region and 2 Midline of the Central Brain (CB). The demarcated ROIs are indicated in Figures 4i and S5r, and the equivalent ROI were used for Ubq signal intensity quantifications in all samples analysed (Figure 4h-k and S5q-r). The 1000μm^2^ (31.62μm x 31.62μm) ROI areas quantified represent the average pixel number in this region per μm^2^ and for all genotypes quantified 12 brain hemispheres were analysed and an average value was acquired.

### A11 amyloid aggregate surface area quantifications

Quantifications of A11 positive amyloid aggregate surface area were acquired utilising Image J (Figure 5j). The threshold function was utilized to first delineate the presence of amyloid aggregates in the specified ROIs and the same threshold intensity of 107 was used to quantify all samples. The demarcated ROIs are indicated in Figures 5d-e and S6e-f and i-j, and the equivalent ROI of 43.14μm x 91.96μm were used for A11 amyloid aggregate surface area quantifications in all samples analysed, within a 5μm Z-projection stack. For all genotypes quantified a minimum of 12 brain hemispheres were analysed within the selected ROI region (43.14μm x 91.96μm), surrounding the giant glial cells of the optic chiasm.

### DAPI positive Cell counting

DAPI positively stained cells in the entire ROI (43.14μm x 91.96μm), were counted with the Fiji cell counting tool in a 5μm Z-projection stack for all samples (Figure 5f”,g”,h” and I”, quantifications Figure 5k). A total of 12 samples, from 12 individual adult brains were quantified for each genotype.

### Statistical analysis

Statistical analysis was carried out using prism version 7.02 for windows, graphpad software, (La Jolla, California USA, www.graphpad.com). For lifespan analysis the log rank test was utilized, to compare median lifespans of two specified data sets. In the case of survivals assays (Figure 3b) where error bars (SEM) overlapped between two data sets (Figure 3b) cox-regression (cr) analysis was used to calculate statistical significance in these instances. For climbing assays and DA neuron volume quantification, two ways analysis of variance (ANOVA) was used to compare the specified data points (***, p < 0.001; **, p < 0.01 and *, p < 0.05), whereas for quantification of DA neuron number, Ubq signal intensity, and A11 positive amyloid aggregates, the two-tailed t-test was used.

Analysis of Real Time PCR were performed as follows: i) when comparing 2 samples, equal variances were confirmed by an F test. ii) when comparing multiple samples, equal variances were confirmed by Bartlett and Brown-Forsythe tests. In all cases normality of data was assessed with Shapiro & Wilk test and parametric or non-parametric tests were used accordingly. Since all data passed normality test, significance was determined by two-tailed T-test or by One-way ANOVA with *post hoc* Tukey Multiple Comparison Test (***, P < 0.001; **, P < 0.01 and *, P < 0.05). Statistical analysis was carried out using Prism version 7.02 for Windows, GraphPad Software, La Jolla California USA, www.graphpad.com

## Acknowledgments

We thank all ‘‘fly’’ members of the IBV Institute, and Tamas Matusek, Laurence Staccini-Lavenant and Caterina Novelli for additional help. We also thank Julien Marcetteau, Pierre Léopold, Florence Besse, Caroline Médioni, Arnaud Hubstenberger, Hervé Tricoire, Thomas Préat, Pauline Spéder, Laure Bally-Cuif and Catherine Rabouille for critical analysis of the manuscript. AR held a Ligue Nationale Contre le Cancer fellowship. This work was supported by grants from Ligue National and Contre le Cancer ‘‘Equipe labellisée 2016’, the LABEX SIGNALIFE (ANR-11-LABX-0028-01), the ANR (ANR-15-CE13-0002-01 and ANR-17-C814-0041-01) to P.P.T, the Federation de la Recherche sur le Cerveau to S.B.

## Authors contributions

A. R. and P. P. T. conceived the experiments and wrote the manuscript. J. N. and M. R. conducted the experiments on Figures 1a-b, 3n, 4a-d, 4m-n S1f-g’’’, S5a-d and S5t-u. S.B. and AH conducted experiment on Figure 3o. A. R. conducted all other experiments. All authors participated in the critical analysis of the manuscript.

---

**Fig. S1.**
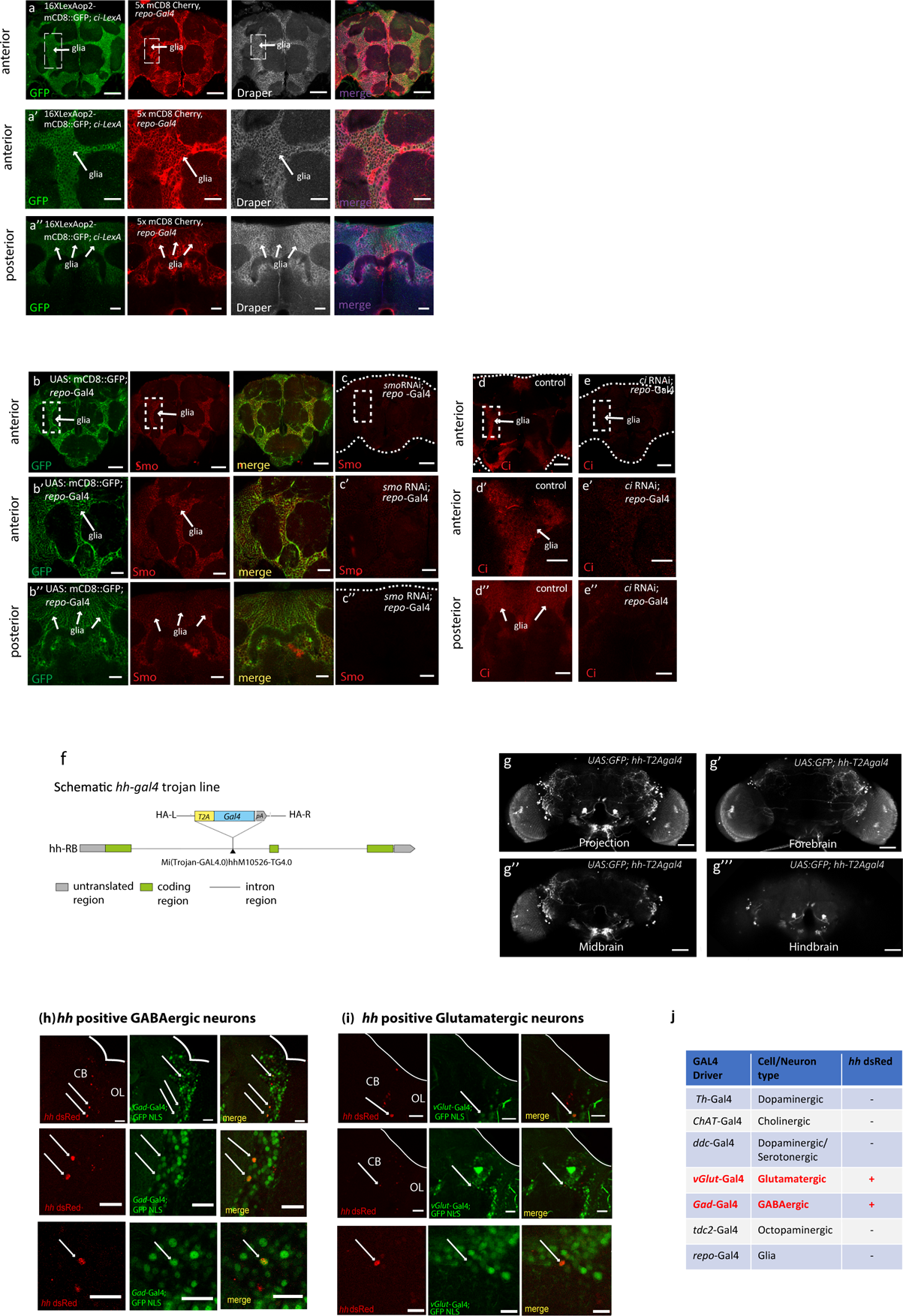
Location and expression of Hh signalling components in the adult brain, **Related to** Figure 1. (a-a’’) *ci*-expressing cells have glial identity (a) *ci*-*lexA* driving mCD8::GFP (green) co-labelled with the glial markers *repo*-Gal4 5xmCD8 Cherry (red) and Draper (grey). Scale bars: 50μm. Equivalent panels in a’ represent 2x magnified images of the anterior (1μm) section of the adult brain, demarcated by the boxed region. (a’’) The equivalent posterior section of the brain is shown in the lower panels. (b-e) Smo/Ci immunoreactivity is reduced upon RNAi knockdown of *smo* (c-c”) and *ci* in the glia (e-e”), compare to controls in b and d, scale bars 20μm. Magnification of the anterior (1μm) section of the brain (region outlined in upper panel) is shown in middle panel, scale bars 50μm. Posterior (1μm) sections of control brains stained with anti-Smo (b”) and anti-Ci (d”), and the corresponding RNAi knockdown are shown in b” and d”. The left panels in a-a” show labelling of the glia using mCD8::GFP and the merge with the Smo immunostaining is shown in the right panels. (f) Schematic representation of the *hh*-T2A-Gal4 trojan line. Projection pattern for *hh* expressing cells. (g-g’’’) Cells positive for *hh-*T2A*-*Gal4, expresses cytoplasmic GFP, (g-g’’’) images of 1μm sections of the forebrain, midbrain and hindbrain respectively). (h-i) *hh* neurons are GABAergic and glutamatergic. The *hh*-dsRed enhancer trap (which labels nuclei of *hh*-expressing cells – left panels) overlaps (white arrows) with a nuclear GFP driven by either GAD-Gal4 or vGlut-Gal4 (middle panels – green channels). Genotypes: UAS: GFP NLS, *hh*-dsred/GAD-Gal4 and UAS: GFP NLS, *hh*-dsRed/vGlut-Gal4. Middle panels (h-i) and lower panels (i) are magnified images of single GABAergic and Glutamatergic cell body neurons positive for *hh-*dsRed (white arrows). Scale bars: 20μm and magnified images 5μm and 10μm. CB and OL stand for central brain and optical lobe, respectively (Images consist of a 1 μm section). (j) Table showing the subtypes of neuronal cells tested for overlaps with *hh* positive cells.

**Fig S2.**
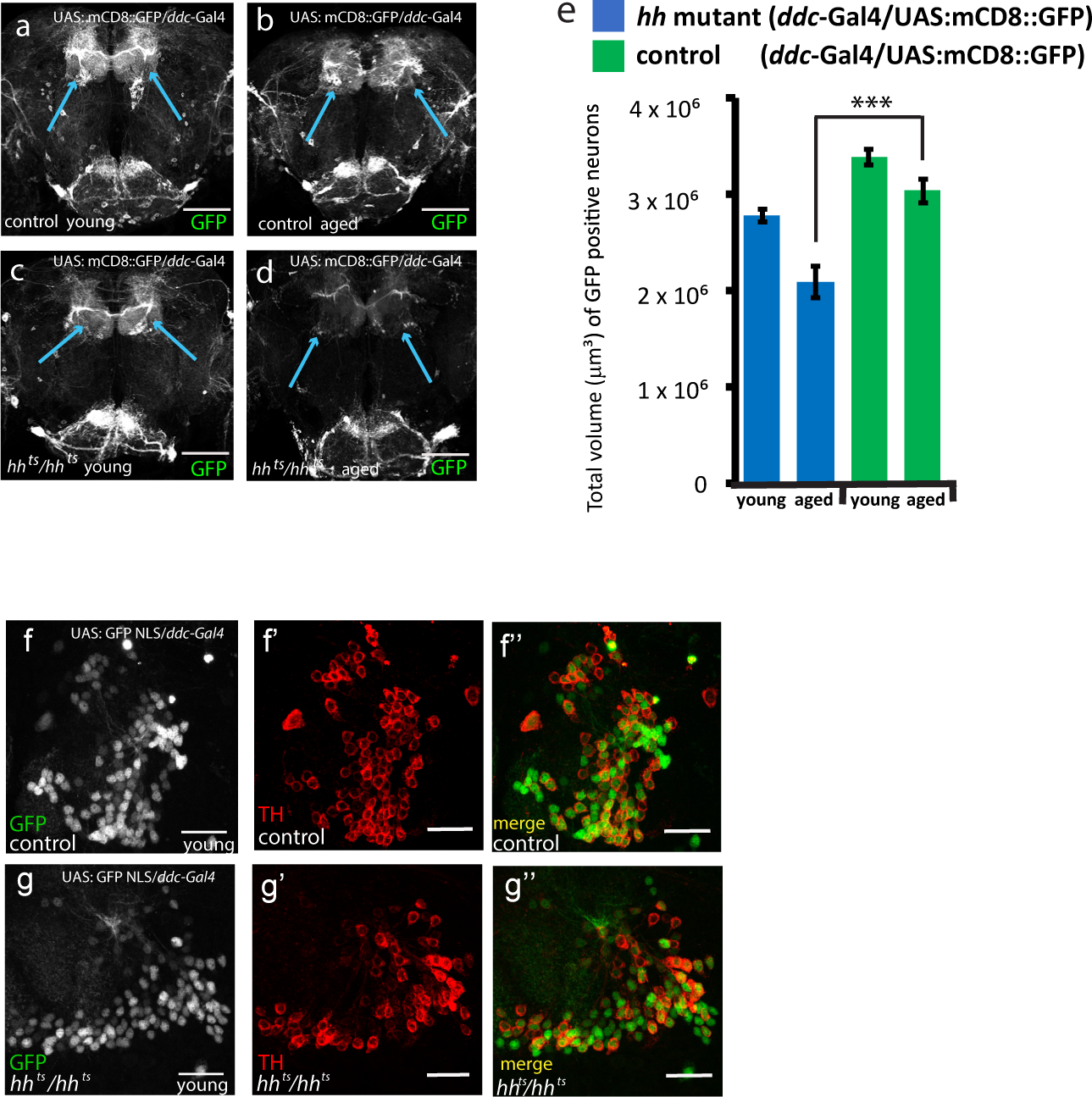
DA neuron loss in hh mutant flies, Related to Figure 1. (a-d) Representative images of *hh* mutant and control adult fly brains with DA neurons labelled with mCD8::GFP, in young (a, c) and aged flies (b, d). Arrows label the PAM DA neurons. Scale bars: 50μm, Z-projection stacks are of the entire adult brain. (e) Quantifications of DA neuron volume in the control and *hh* mutant adult flies using a mCD8::GFP membrane marker. (f-g’’) Representative confocal images of the PAM cluster cell bodies of control (f-f’’) and *hh* mutant (g-g’’) in young adult fly brains, in which DA neurons are labelled with a nuclear GFP marker (GFP NLS in grey) and with anti-TH immunostaining (in red), Scale bars:10μm. Quantifications in Fig. 1i.). For DA neuron number and DA neuron volume quantification two-way ANOVA test was used to compare the specified data points for *hh* mutant and control samples (***, p < 0.001).

**Fig. S3.**
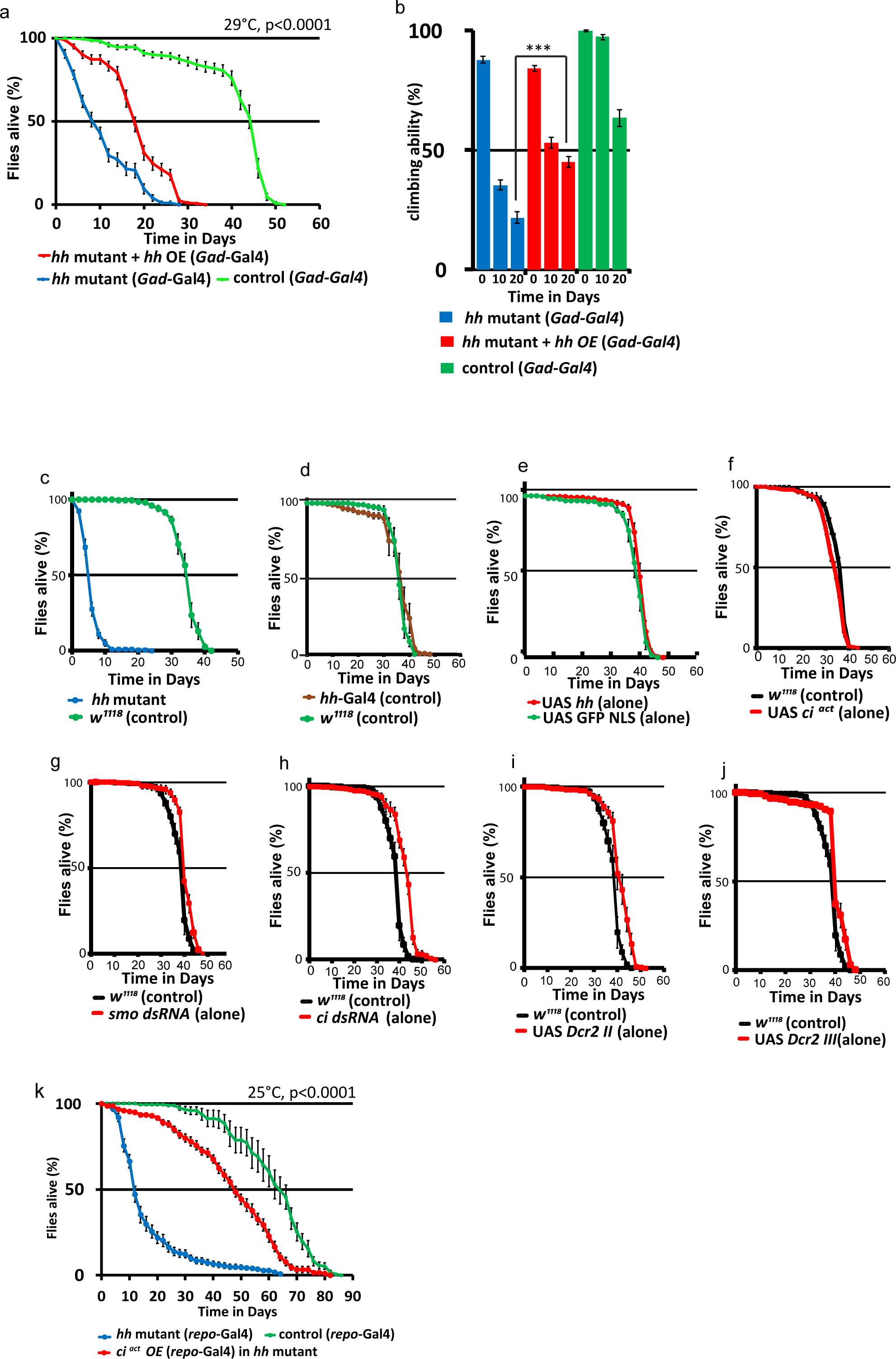
Hh signalling in GABAergic neurons partially rescues the *hh* mutant shortened lifespan and defective locomotion, Related to Figure 2. (a) Overexpression of *hh* in GABAergic neurons partially rescues the *hh* mutant shortened survival phenotype and (b) the impaired locomotion phenotype. (b) *hh* mutant male flies (*hh^ts2^/hh^ts2^*) exhibit a reduced survival time. (d-j) Survival time of UAS and Gal4 control lines: (d) *hh*-Gal4, Gal80^ts^/+; (e) UAS:*hh*M4; (f) UAS:*ci^act^*/+; (g) UAS:*smo* dsRNA/+; (h) UAS:*ci* dsRNA/+; (i) UAS:Dcr2 (II)/+; (j) UAS:Dcr2 (III)/+. All compared to control *w^1118^* (c, d, f-j) or UAS-GFP NLS flies (e). (k) Expression of an activated form of Ci in the glia (*ci^act^* OE, *repo*-Gal4) rescues the short survival time of *hh* mutant at 25°C. For lifespan analysis log rank test was utilised (p<0.0001), for climbing activity two-way anova test was used, p<0.001***.

**Fig S4.**
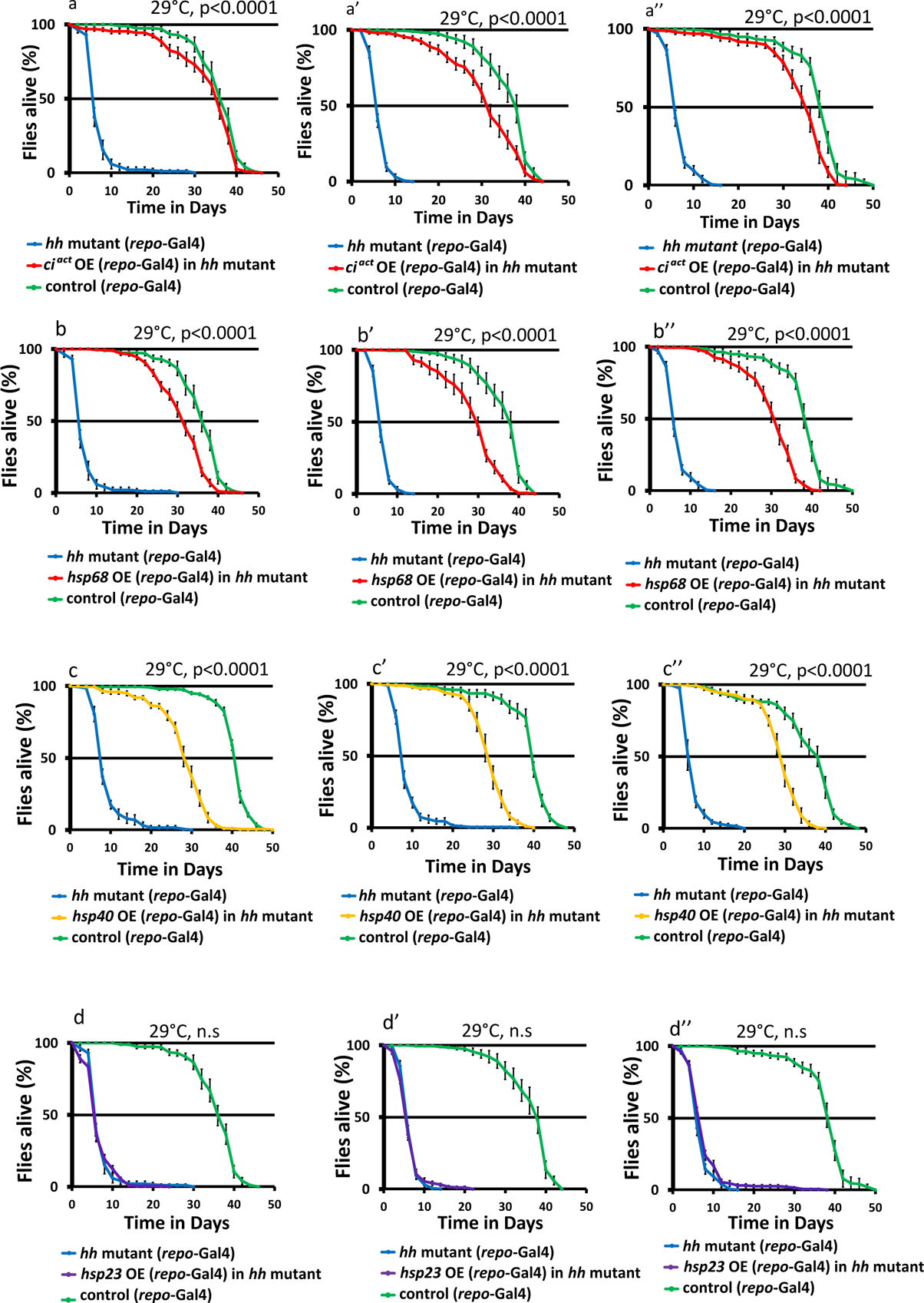
Stimulation of Hh signalling and expression of Hh target chaperones in the glia potently rescues the *hh* mutant shortened lifespan, Related to Figure 3 and 4. (a-a”) Three independent experiments performed in triplicate, showing that overexpression of *ci ^act^* rescues the *hh* mutant short-lived phenotype. Overexpression of *hsp68* (b-b’’) and *hsp40* (c-c”) potently rescue the *hh* mutant short-lived phenotype, (log rank test – p<0.0001) whereas no rescue was observed with overexpression of *hsp23* (log rank test – ns) (d-d”). Note that longevity curves for *hsp68* and *hsp23* were run in parallel with the same controls. For lifespan analysis log rank test was utilised and *hh* mutant samples were compared to *hh* mutants expressing the specified chaperone in the glia, see Table S2 and Table S5.

**Fig. S5.**
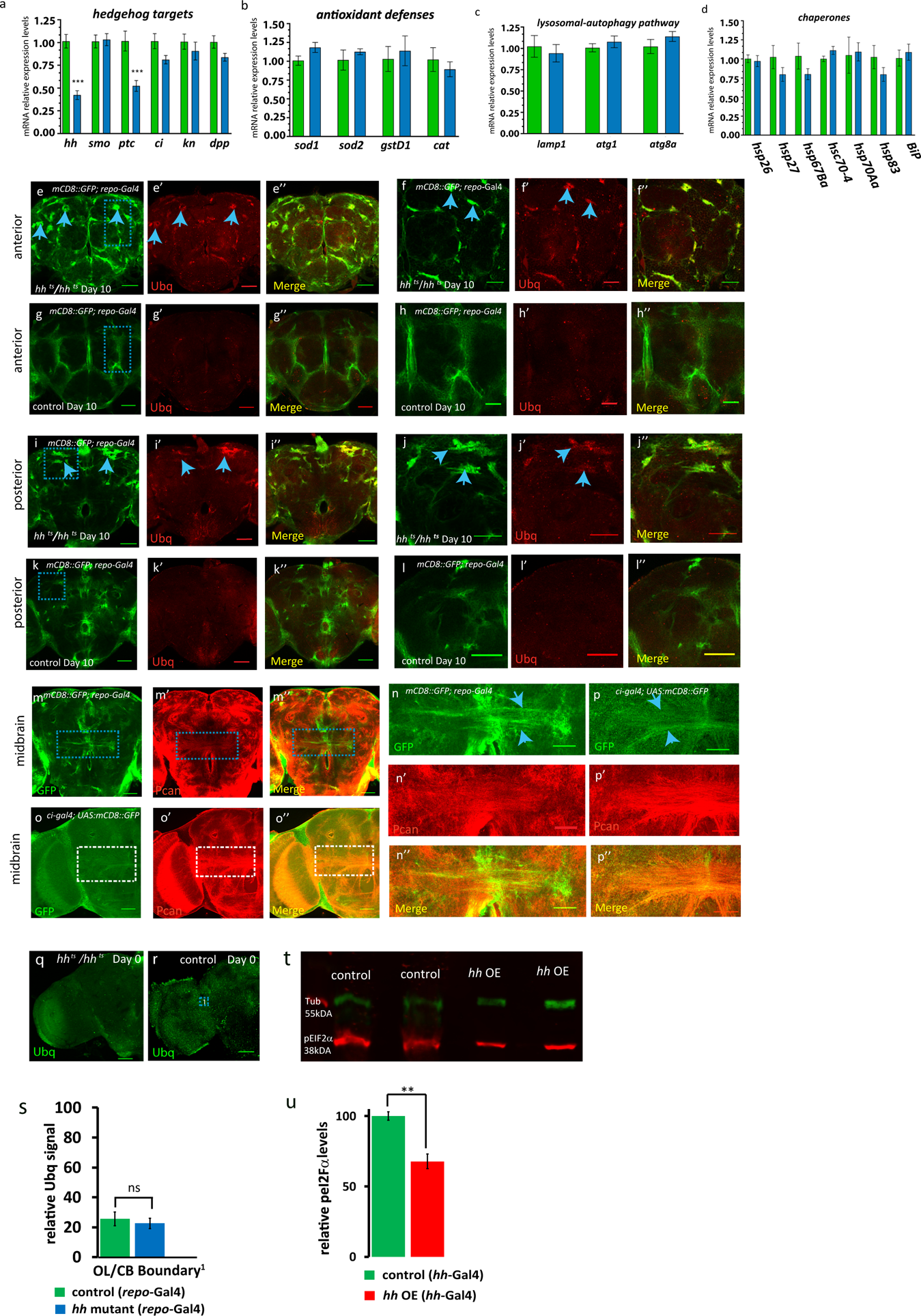
Proteostasis defects in *hh* mutant flies, Related to Figure 4. (a) mRNA expression level of Hh signalling pathway components in adult heads. Both *hh* and *ptc* mRNA levels were substantially decreased in *hh* mutant samples compared with age-matched controls (p<0.001***, two tailed t-test) (b-d) Among several tested genes, the mRNA levels of genes implicated in antioxidant defences (b), the lysosomal autophagy pathway (c) and chaperones (d) exhibit no significant difference except for the ones presented in figure 4b. Each experiment consisted of 4-5 independent biological replicates. The genes and the pairs of primers used for the analysis are listed in Methods. (e-l”) Representative images of day 10 *hh* mutant (n=10) and control fly brains (n=10) with glial cells labelled with mCD8::GFP (green channel). In all analysed *hh* mutant samples we observed abnormal accumulations of GFP in the glia, which also contain elevated level of ubiquitin (Ubq) staining, (indicated by blue arrows – red channel) in both anterior (e-e”) and posterior brain regions (i-i”). This phenomenon was not observed in the equivalent aged matched control samples in either the anterior (g-g”) or posterior brain regions (k-k”). 1μm section, scale bars 50μm. The equivalent magnified image of the region of interest (marked by the blue dashed rectangle) are displayed in the adjacent panels (f-f”, h-h”, j-j” and l-l”), respectively. 1μm stack, scale bars 30 μm. (m-n”) Control flies with glial cells labelled with mCD8::GFP (green channel) in the midbrain region m), immunostained with anti-Perlecan (Pcan, red in m’), merged channels (m’’). 1μm section, scale bars 50 μm. (n) Magnification of the region of interest (demarcated by the blue dashed rectangle in m) of the glial tracts in the adult midbrain are displayed in the adjacent panels (n-n”). Stack of 2 x 1μm sections, scale bars 50 μm. (o) mCD8-GFP driven in ci-expressing cells immunostained with Pcan (o’) and merged image in o’’, scale bars 50 μm. Magnification of the region of interest (demarcated by the white dashed rectangle in o) are displayed in the adjacent panels (p-p”). Stack of 2 x 1μm sections, scale bars 30 μm. (q-r) Images of Day 0 *hh* mutant and control brains immunostained for Ubiquitin (Ubq, green), scale bars 50 μm, quantifications, (s, see Fig. 4l-l’), two-tailed t-test (ns). (t) Representative Immunoblot showing pEIF2a levels in *hh* overexpressing day 30 adult head samples and aged matched controls. (u) *hh* overexpressing samples exhibit a 32% reduction in pEIF2a levels compared to age-matched control specimens, (p<0.005**, two-tailed t-test).

**Fig. S6.**
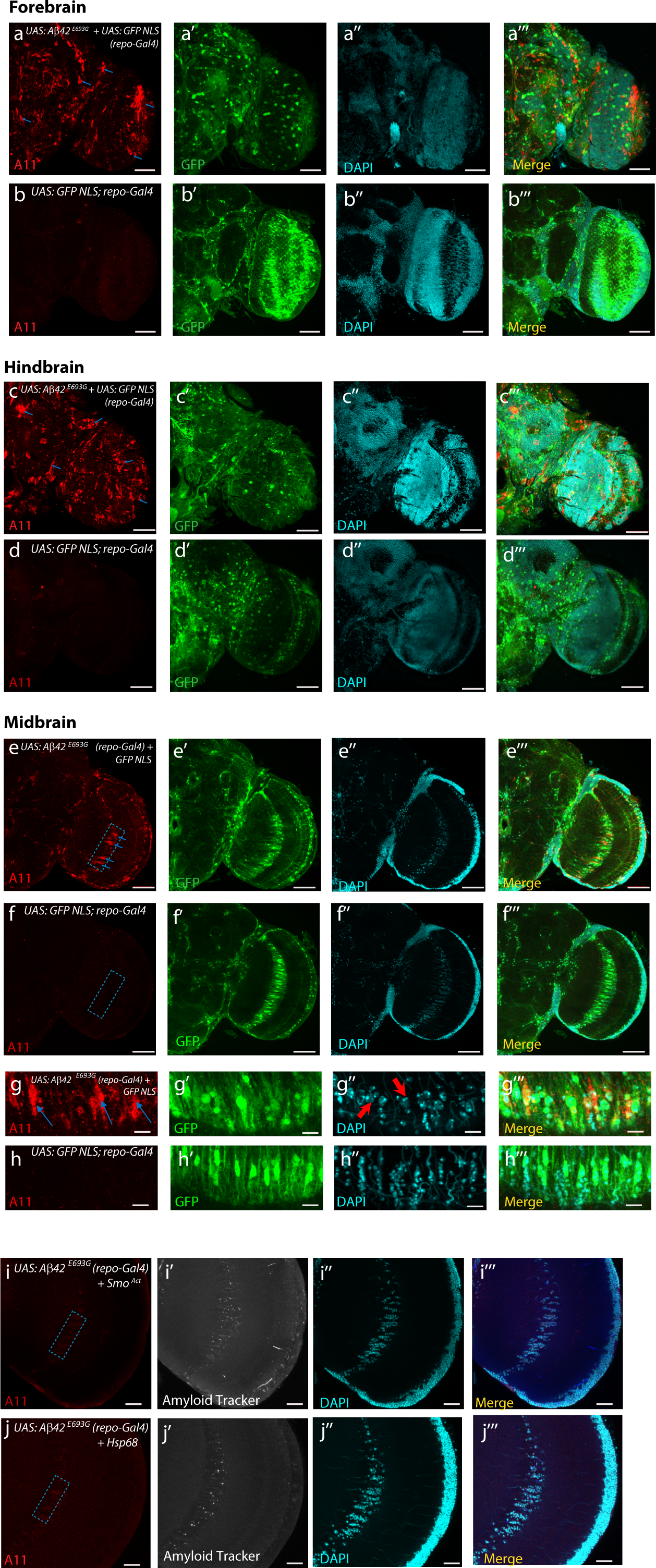
Detection of Amyloid aggregates in adult brains of flies expressing the human Aβ42^E693G^ variant in glial cells, Related to Figure 5. Representative images of 1μm sections for the forebrain (a-a’’’), hindbrain (c-c’’’) and midbrain (e-e’’’, 5μm section) expressing the human Aβ42 ^E693G^ variant along with GFP NLS (green channel) in the glia and stained for DAPI (cyan channel), along with the merged channel in 22 day old specimens. The equivalent age-matched control samples expressing only GFP NLS in the glia are also shown: forebrain (b-b’’’); hindbrain (d-d’’’) and midbrain (f-f’’’) sections. Throughout all sections of the adult brain amyloid aggregates (indicated by blue arrows) can be observed in the samples expressing human Aβ42 ^E693G^ in the glia, whereas this is not the case for the control specimens, immunostained with the A11 amyloid aggregate specific antibody, Scale Bars: 50μm; The demarcated dashed box (ROI) in e, encompasses the giant glial cells of the optic chiasma and the magnified images (g-g’’’) reveal the presence of amyloid aggregates indicated by the blue arrows (g), whereas in the equivalent control specimens (h-h’’’) such aggregated structures could not be detected (h). Samples expressing human Aβ42 ^E693G^ in the glia also exhibited enlarged cells with a swollen morphology (red arrows) and a reduction in cell numbers in the ROI compared to control samples. Scale Bars: 10μm. (i-j) Confocal section (5μm stack) of the optical lobe of 15 day old flies expressing human Aβ42 ^E693G^ along with Smo^Act^ (i) or Hsp68 (j) in the glia, Scale Bars 20μm. In order to quantify the surface area of amyloid aggregates and cell number in the region of interest - ROI (dashed box), an area of 43.14μm x 91.96μm was selected (see Figure 5g-h, for selected ROIs).

**Table S1.** A summary of the DA neuron volume data and statistical analysis for all the relevant genotypes shown in the text. (Statistical comparisons between age-matched samples for young, middle aged and old flies are colour coded blue, green and yellow respectively).

| Genotype | Figure | young | P value | middle age | P value | aged | P value |
| --- | --- | --- | --- | --- | --- | --- | --- |
| **DA neuron network volume of Hh mutants (mCD8::GFP labelling)** |  |  |  |  |  |  |  |
| *hh<sup>ts2</sup>, Ddc-GAL4/ UAS:mCD8::GFP, hh<sup>ts2</sup>* | S2c-d | 2780914 | n=12 (<0.01)** | 2348623 | n=10 (<0.001)*** | 2094783 | n=4 (<0.001)*** |
| control: *Ddc-GAL4/ UAS:mCD8::GFP* | S2a-b | 3391080 | n=10 (<0.01)** | 3068557 | n=10 (<0.001)*** | 3041423 | n=8 (<0.001)*** |
| **DA neuron network volume of Hh mutants (TH immunostaining)** |  |  |  |  |  |  |  |
| *hh<sup>ts2</sup>, Ddc-GAL4/ UAS:mCD8::GFP, hh<sup>ts2</sup>* | 1m, 1n | 1323223 | n=12 (n.s) | 1097554.8 | n=10 (<0.05)* | 651896 | n=4 (<0.001)*** |
| control: *Ddc-GAL4/ UAS:mCD8::GFP* | 1k, 1l | 1331695 | n=10 (n.s) | 1272523.2 | n=10 (<0.05)* | 1178530 | n=8 (<0.001)*** |

| Genotype | Figure | 0d | P value | 20d | P value |
| --- | --- | --- | --- | --- | --- |
| **DA neuron volume of Hh mutant flies expressing Ci Act In glia (TH immunostaining)** |  |  |  |  |  |
| control: *UAS:GFP NLS/GAL80<sup>ts</sup>;Repo-GAL4/+* | 3g,3j | 1152009 | (n.s) n=9 | 995048.5 | n=6 (<0.001)*** |
| *GAL80<sup>ts</sup> /+; UAS: GFP NLS, hh<sup>ts2</sup> /Repo-GAL4, hh<sup>ts2</sup>* | 3h, 3k | 1135597 | n=10 | 341226.125 | n=8 |
| *GAL80<sup>ts</sup> /+; UAS: Ci PKA, hh<sup>ts2</sup> /Repo-GAL4, hh<sup>ts2</sup>* | 3i, 3l | 1166615 | (n.s) n=10 | 698677.4 | n=5 (<0.001)*** |

**Table S2.**
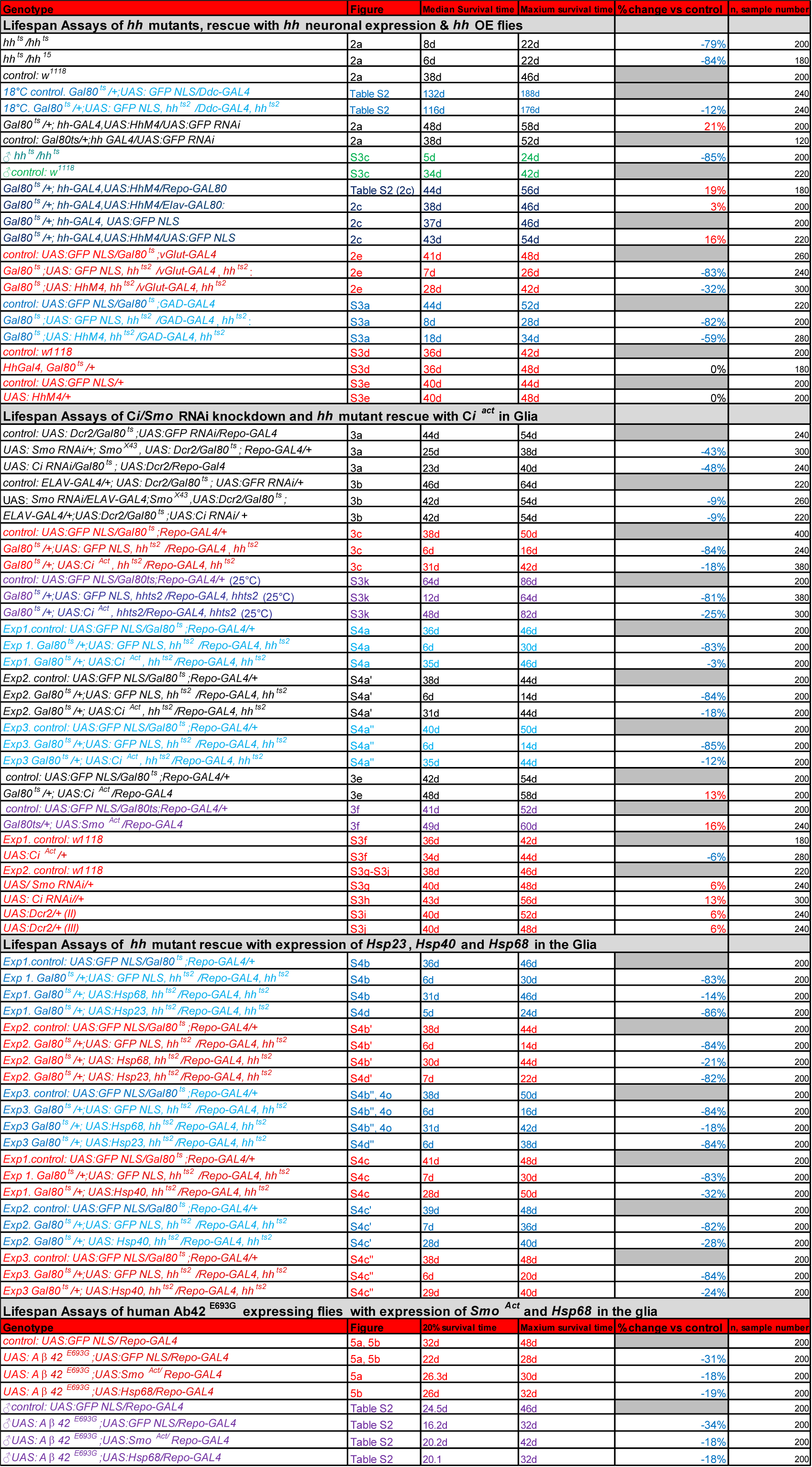
Summary of statistical analysis/median and maximum lifespan values for all relevant genotypes mentioned in the text.

**Table S3.** A summary of the climbing ability and a statistical analysis for all the relevant genotypes mentioned in the text.

| Genotype | Figure | 0 Day climbing ability | Aged climbing ability | Two-way Anova test |
| --- | --- | --- | --- | --- |
| **Climbing ability of *hh* mutants, rescue with *Hh* neuronal expression & *Hh* OE flies** |  |  |  |  |
| control: *w<sup>1118</sup>* | 2b | 93±0.8%; 83±1.4% (5d) | 66±2.2% (15d) |  |
| *hh<sup>ts2</sup>/hh<sup>ts2</sup>* | 2b | 86±1.8%; 30±2.7 (5d) | 4±1.1% (15d) | <0.001 *** |
| *Gal80<sup>ts</sup>/+; hh-GAL4/UAS:HhM4* | 2b | 97±0.8%; 95±1.0 (5d) | 88±1.6% (15d) | <0.001 *** |
| *Gal80<sup>ts</sup>/+; hh-GAL4,UAS:HhM4/Elav-GAL80 (8cm)* | 2d | 96±1.1% | 20±1.5% (35d) | <0.001 *** |
| *Gal80<sup>ts</sup>/+; hh-GAL4,UAS:HhM4/+ (8cm)* | 2d | 96±1.0% | 39±1.7% (35d) |  |
| *Gal80<sup>ts</sup>/+; hh-GAL4/UAS:GFP NLS (8cm)* | 2d | 90±1.7% | 19±2.1% (35d) | <0.001 *** |
| control: *UAS:GFP NLS/Gal80<sup>ts</sup>;vGlut-GAL4* | 2f | 100±0% | 70±2.6% (20d) | <0.001 *** |
| *Gal80<sup>ts</sup>;UAS: GFP NLS, hh<sup>ts2</sup> /vGlut-GAL4, hh<sup>ts2</sup>* | 2f | 84±1.8% | 26±2.3% (20d) | <0.001 *** |
| *Gal80<sup>ts</sup>;UAS: HhM4, hh<sup>ts2</sup> /vGlut-GAL4, hh<sup>ts2</sup>* | 2f | 93±1.3% | 55±2.0% (20d) |  |
| control: *UAS:GFP NLS/Gal80<sup>ts</sup>;GAD-GAL4* | S3b | 100±0.3% | 63±3.4% (20d) | <0.001 *** |
| *Gal80<sup>ts</sup>;UAS: GFP NLS, hh<sup>ts2</sup> /GAD-GAL4, hh<sup>ts2</sup>* | S3b | 88±1.5% | 22±2.4% (20d) |  |
| *Gal80<sup>ts</sup>;UAS: HhM4, hh<sup>ts2</sup> /GAD-GAL4, hh<sup>ts2</sup>* | S3b | 84±1.2% | 45±2.2% (20d) | <0.001 *** |
| **Climbing ability of *Hh* mutant flies expressing *Ci<sup>Act</sup>* in glia** |  |  |  |  |
| control: *UAS:GFP NLS/Gal80<sup>ts</sup>;Repo-GAL4/+* | 3d | 100±0% | 77±2.2% (20d) | <0.001 *** |
| *Gal80<sup>ts</sup>/+;UAS: GFP NLS, hh<sup>ts2</sup> /Repo-GAL4, hh<sup>ts2</sup>* | 3d | 89±1.4% | 17±1.8% (20d) |  |
| *Gal80<sup>ts</sup>/+; UAS: Ci<sup>Act</sup>, hh<sup>ts2</sup> /Repo-GAL4, hh<sup>ts2</sup>* | 3d | 89±1.2% | 54±2.9% (20d) | <0.001 *** |
| **Climbing ability of *Hh* mutant flies expressing *Hsp68* in the glia** |  |  |  |  |
| control: *UAS:GFP NLS/Gal80<sup>ts</sup>; Repo-Gal4/+* | 4p | 94 ±1.2% | 66 ±1.8% (15d) | <0.001 *** |
| *Gal80<sup>ts</sup>;UAS:GFP NLS, hh<sup>ts2</sup> /Repo-GAL4, hh<sup>ts2</sup>* | 4p | 82 ±1.0% | 10 ±1.6% (15d) |  |
| *Gal80<sup>ts</sup>/+;UAS:hsp68, hh<sup>ts2</sup> /Repo-GAL4, hh<sup>ts2</sup>* | 4p | 80 ±1.7% | 38 ± 1.5% (15d) | <0.001 *** |
| **Climbing ability of human Ab42<sup>E693G</sup> expressing flies with expression of *Smo<sup>Act</sup>* and *Hsp68* in the glia** |  |  |  |  |
| control: *UAS:GFP NLS/Repo-GAL4* | 5c | 95±1.0% | 73±2.3% (14d) | <0.001 *** |
| *UAS: A β 42<sup>E693G</sup>;UAS:GFP NLS/Repo-GAL4* | 5c | 82±1.9% | 21±2.7% (14d) |  |
| *UAS: A β 42<sup>E693G</sup>;UAS:Smo<sup>Act</sup>/Repo-GAL4* | 5c | 89±1.3% | 56±1.8% (14d) | <0.001 *** |
| *UAS: A β 42<sup>E693G</sup>;UAS:Hsp68/Repo-GAL4* | 5c | 77±1.2% | 66±1.6% (14d) | <0.001 *** |

**Table S4.**
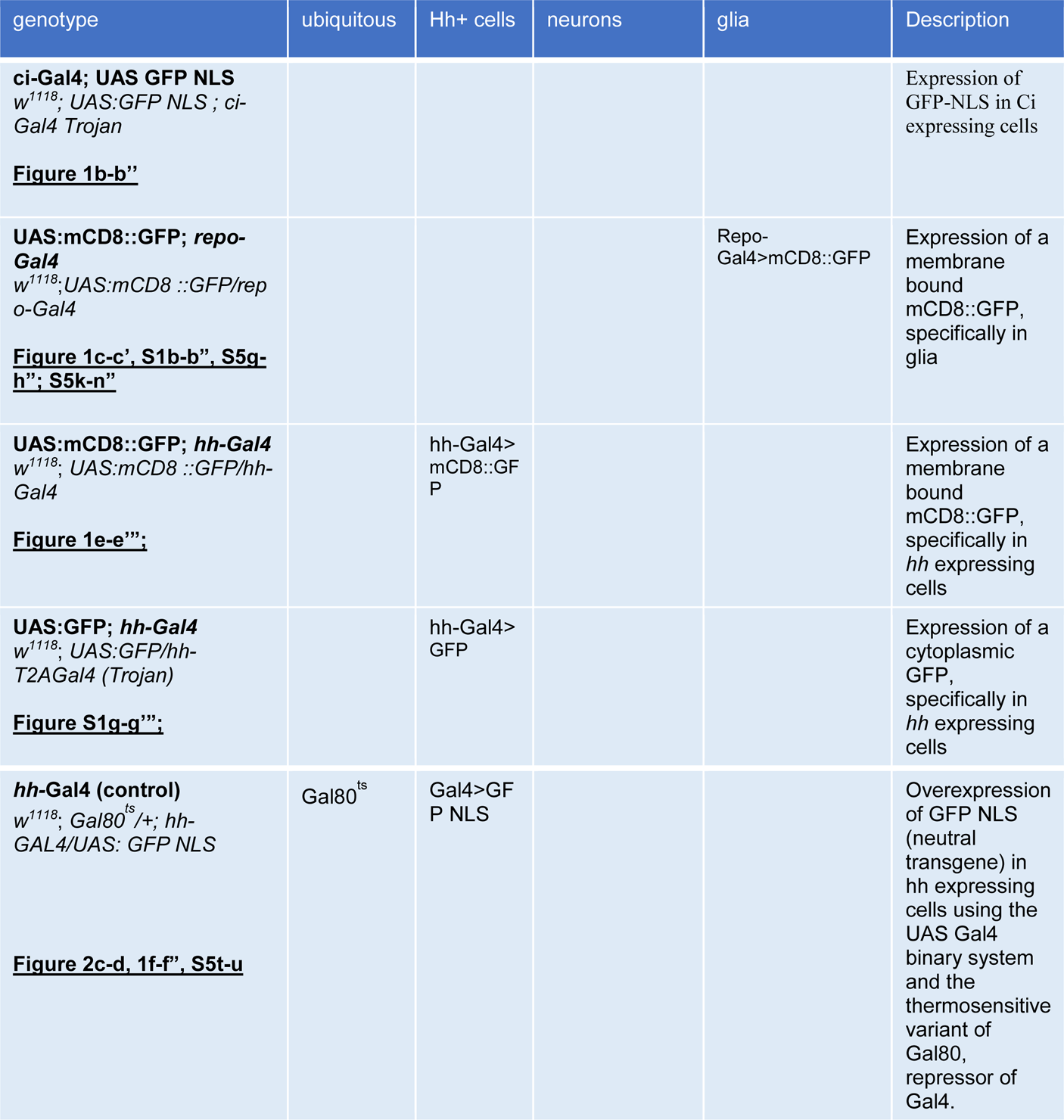
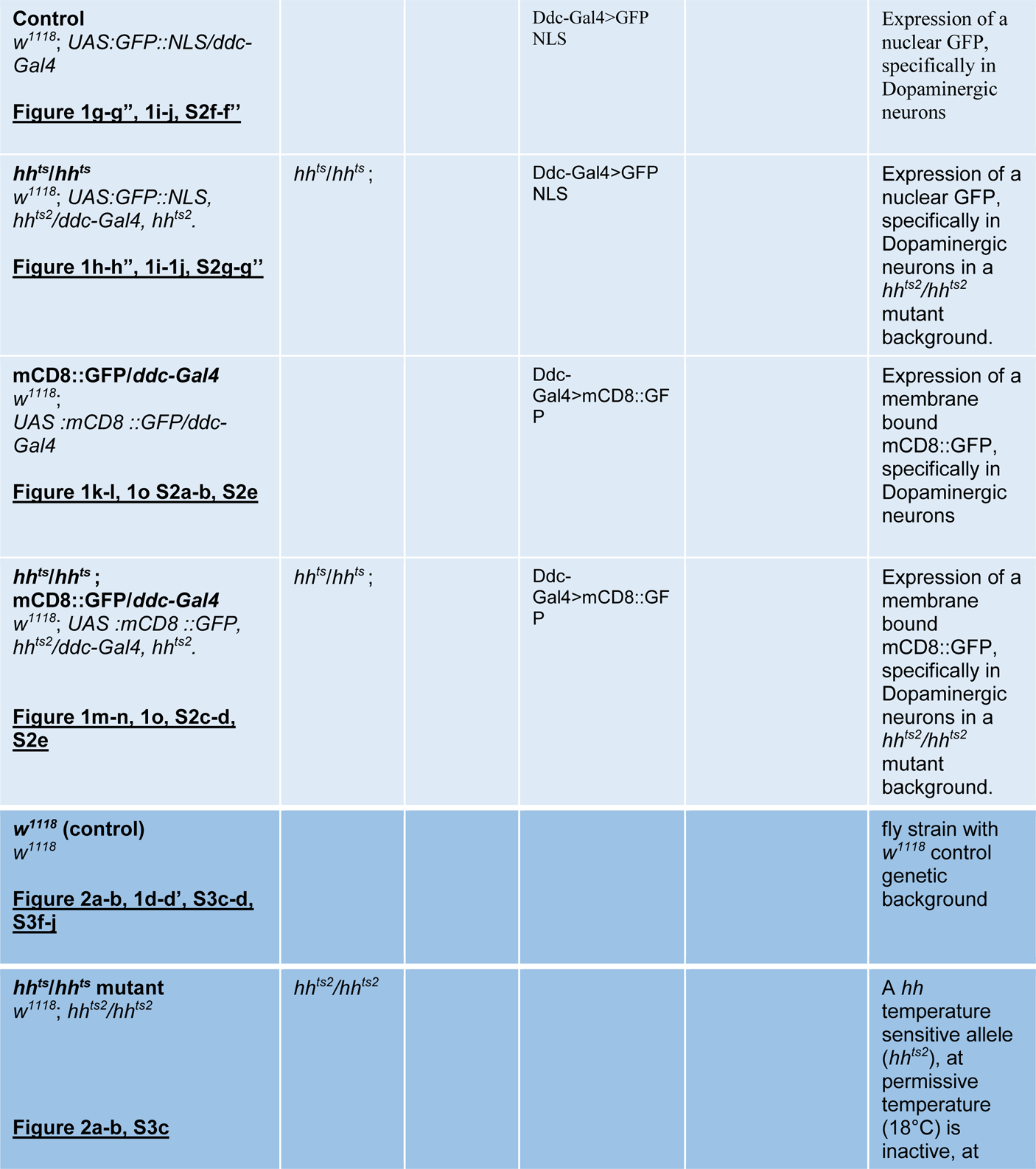
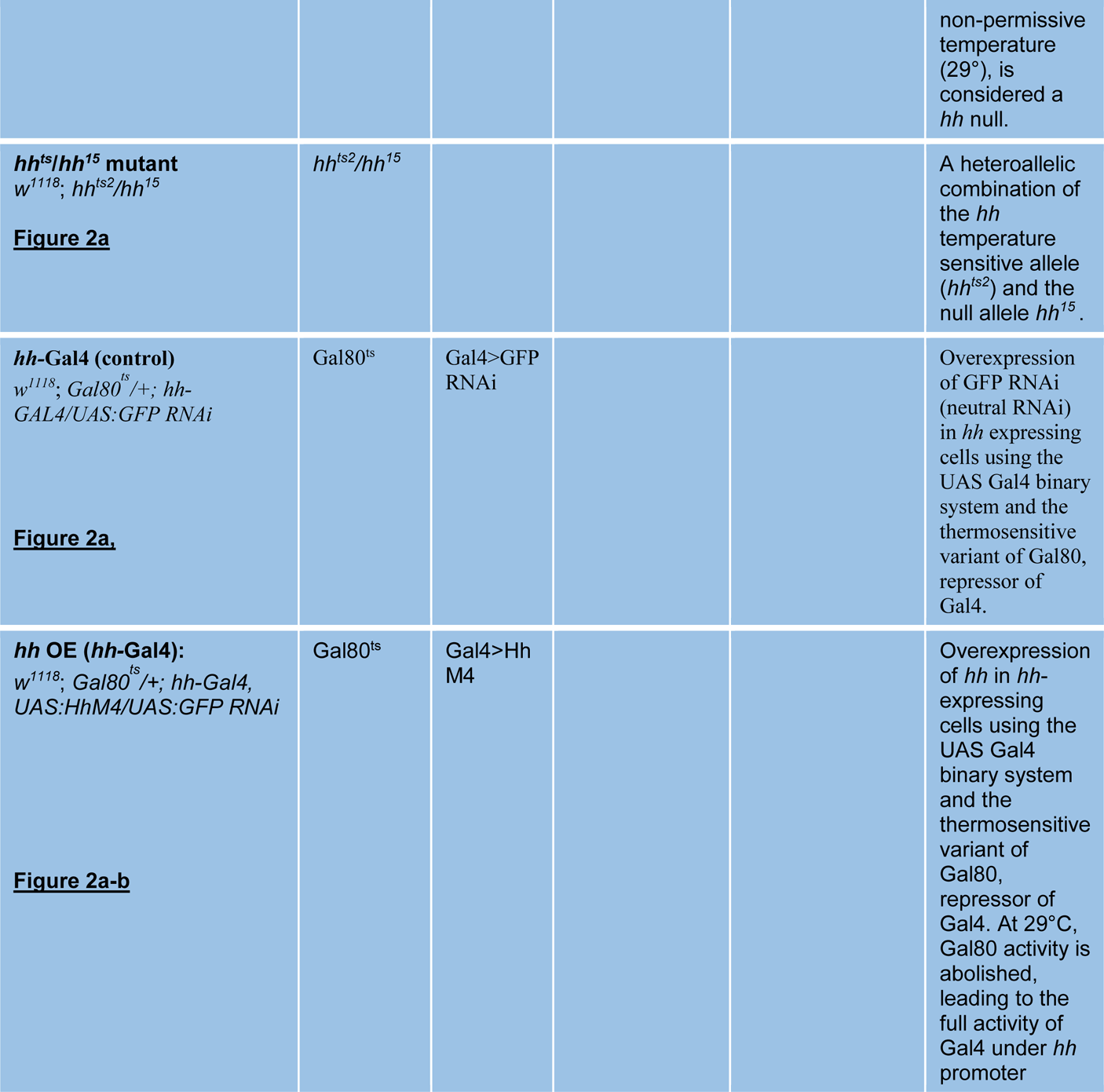
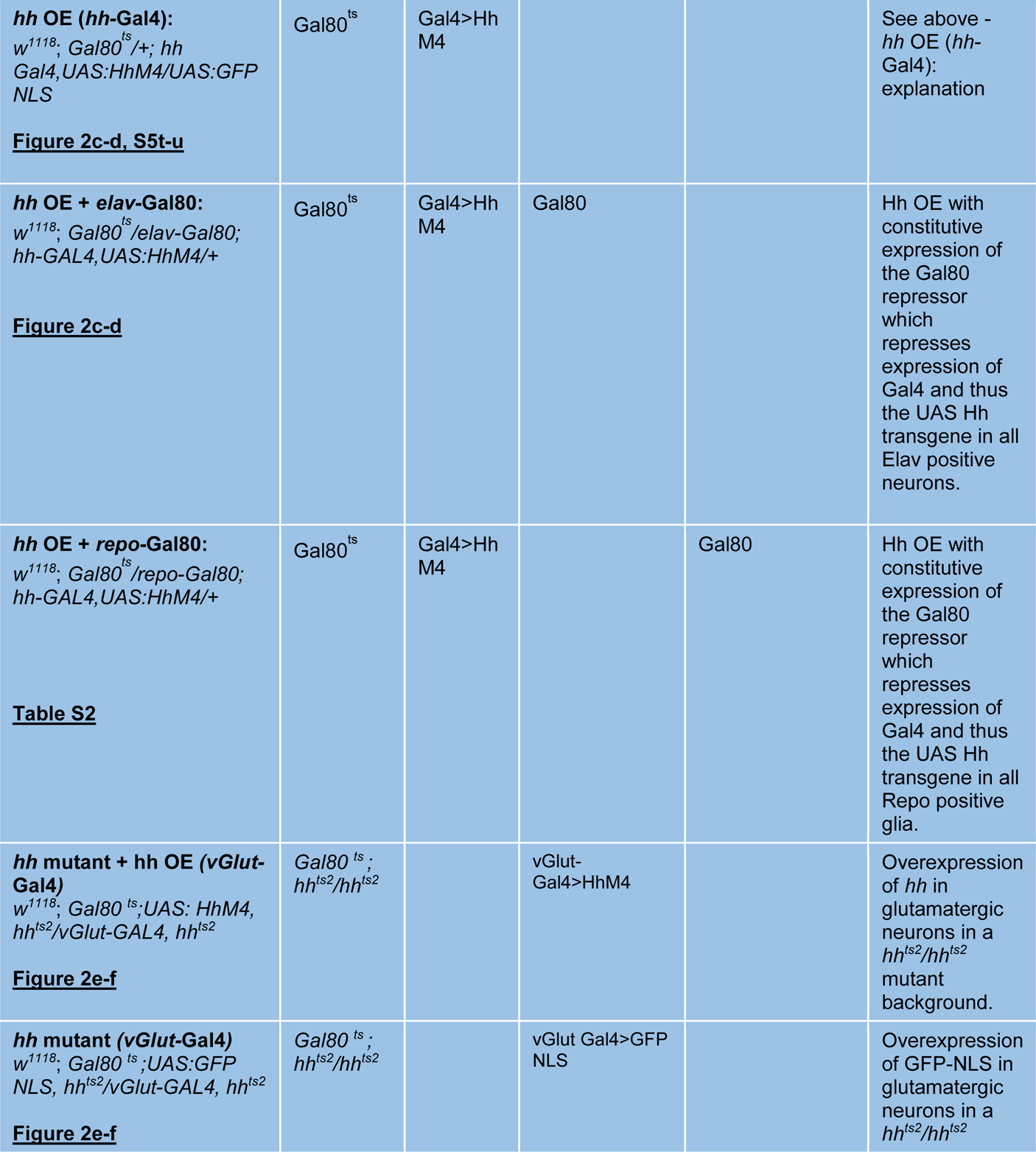
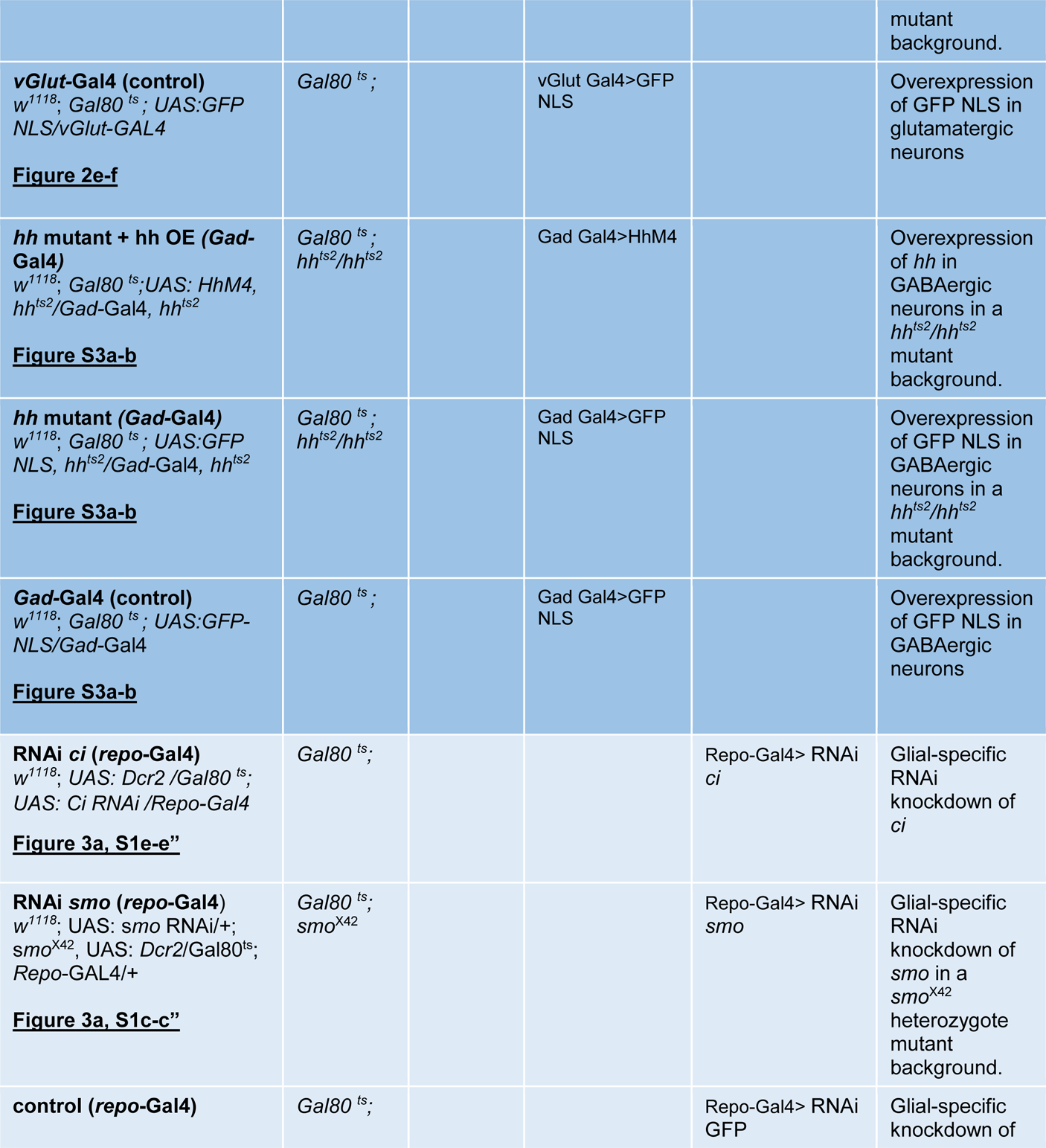
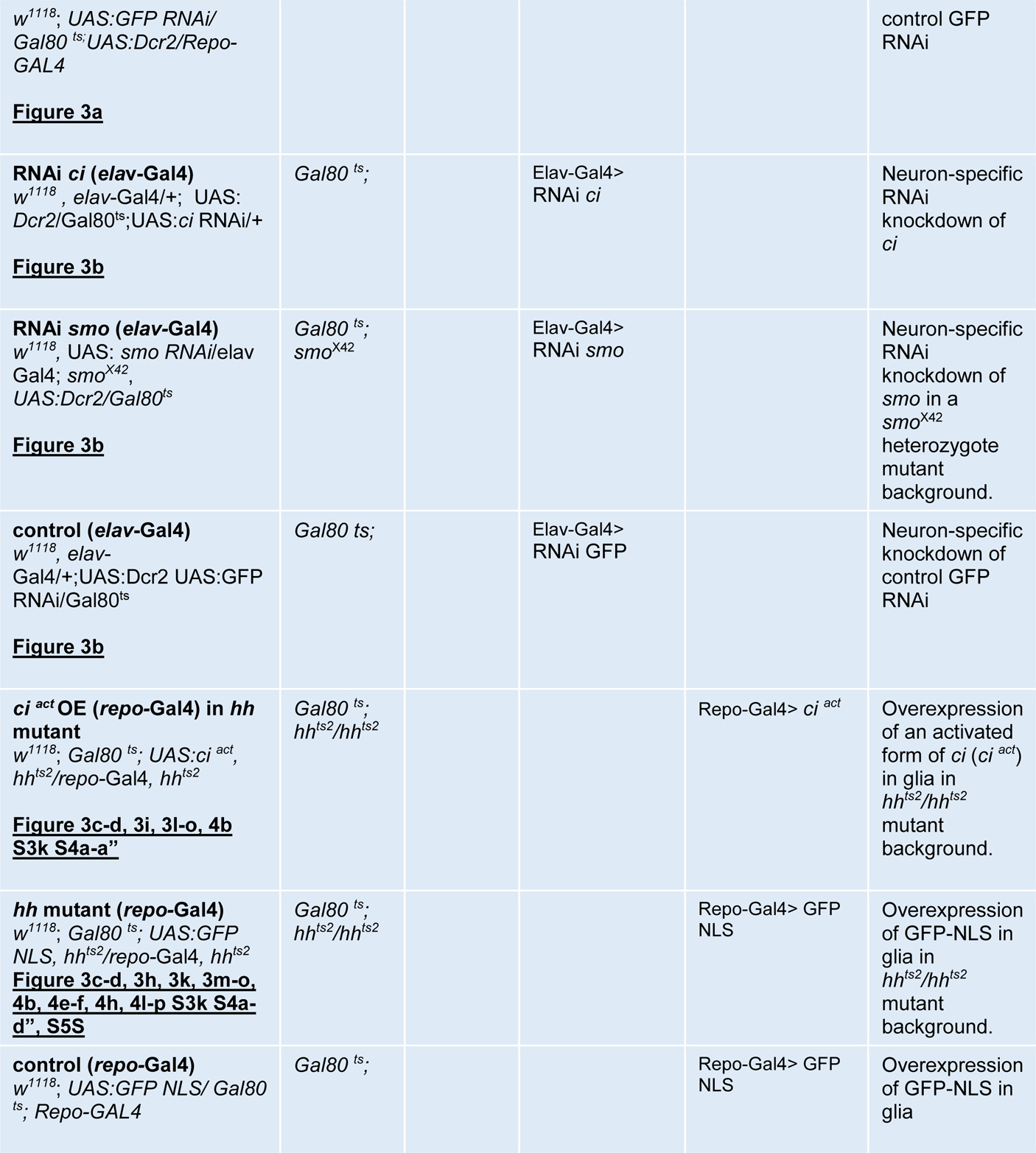
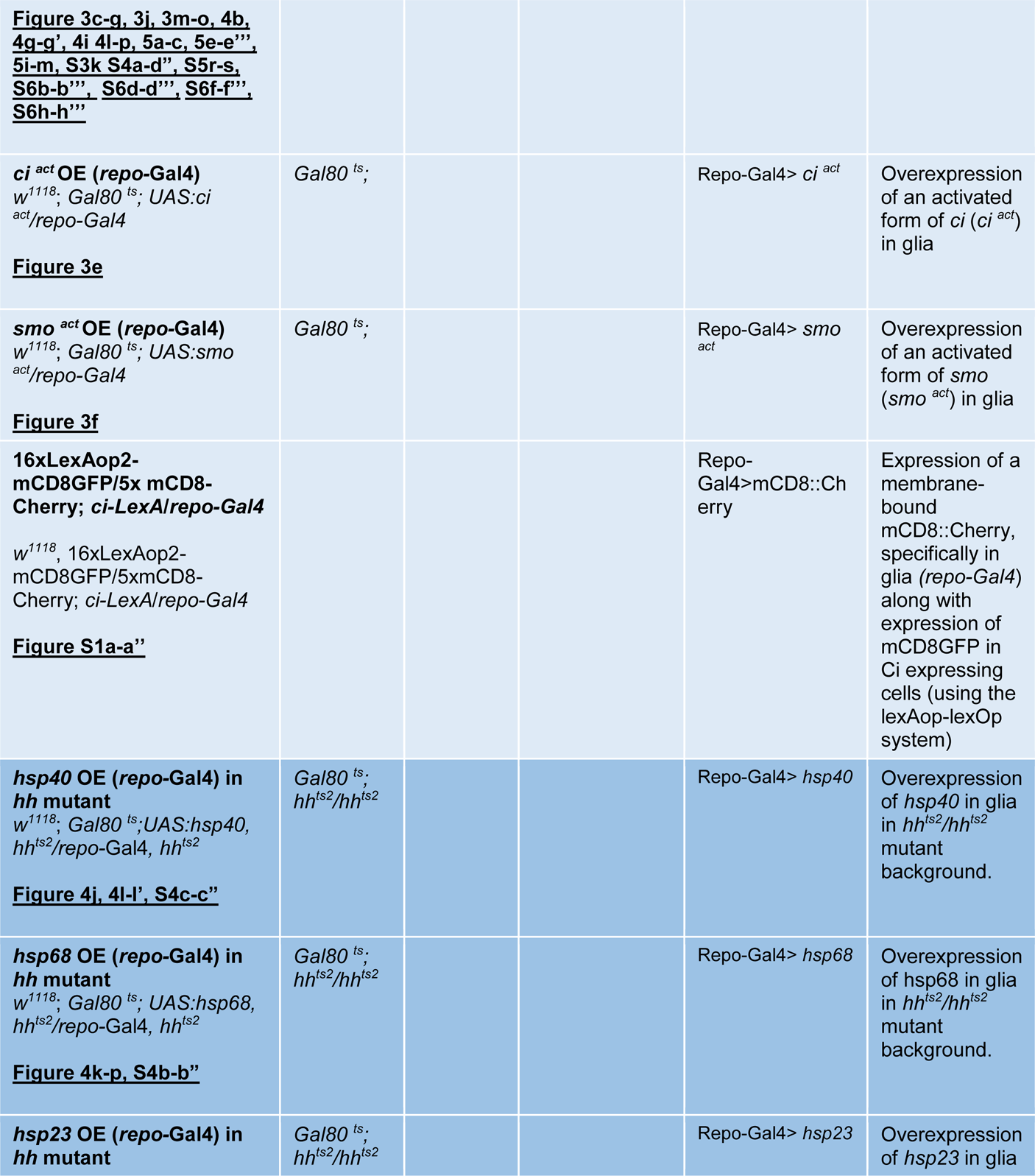
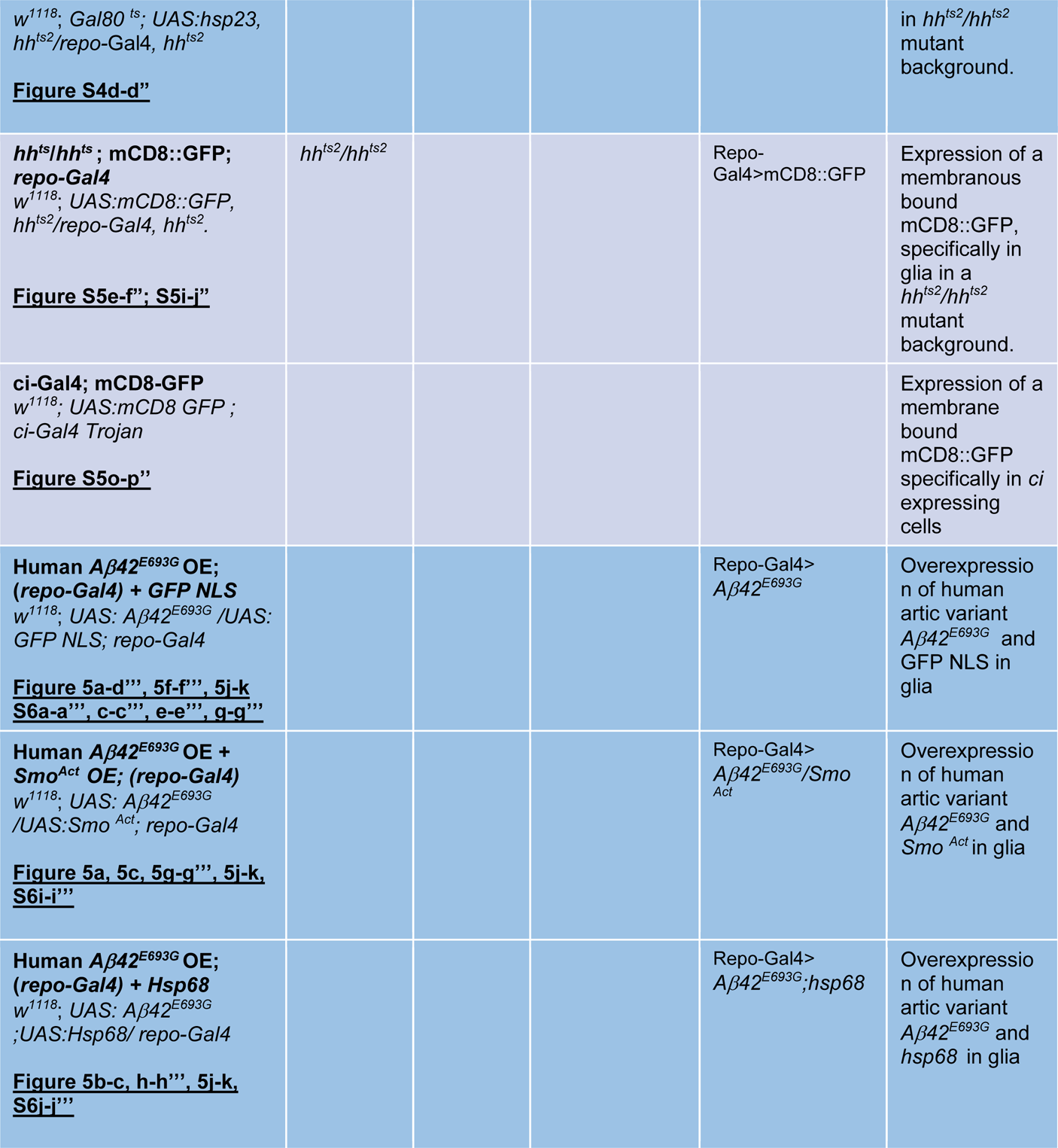
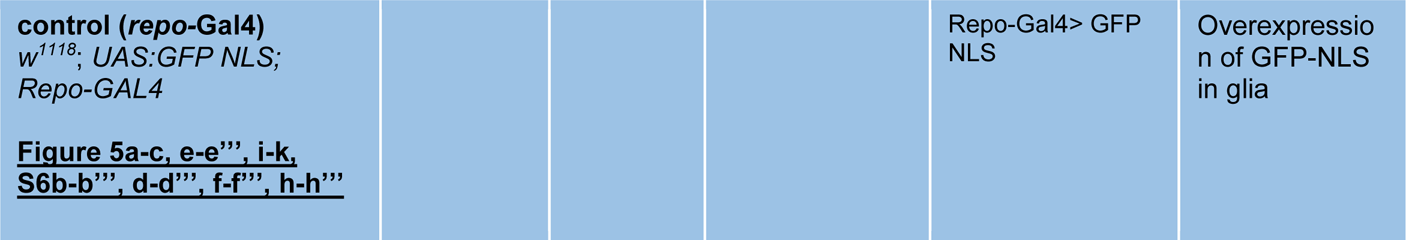
Displays the short-hand notation and the corresponding full genotype used in both the lifespan and locomotion analysis.

| genotype | ubiquitous | Hh+ cells | neurons | glia | Description |
| --- | --- | --- | --- | --- | --- |
| **ci-Gal4; UAS GFP NLS**<br>*w<sup>1118</sup>; UAS:GFP NLS ; ci-Gal4 Trojan*<br>**<u>Figure 1b-b''</u>** |  |  |  |  | Expression of GFP-NLS in Ci expressing cells |
| **UAS:mCD8::GFP; repo-Gal4**<br>*w<sup>1118</sup>; UAS:mCD8 ::GFP/repo-Gal4*<br>**<u>Figure 1c-c', S1b-b'', S5g-h''; S5k-n''</u>** |  |  |  | Repo-Gal4>mCD8::GFP | Expression of a membrane bound mCD8::GFP, specifically in glia |
| **UAS:mCD8::GFP; hh-Gal4**<br>*w<sup>1118</sup>; UAS:mCD8 ::GFP/hh-Gal4*<br>**<u>Figure 1e-e'';</u>** |  | hh-Gal4>mCD8::GFP |  |  | Expression of a membrane bound mCD8::GFP, specifically in *hh* expressing cells |
| **UAS:GFP; hh-Gal4**<br>*w<sup>1118</sup>; UAS:GFP/hh-T2AGal4 (Trojan)*<br>**<u>Figure S1g-g'';</u>** |  | hh-Gal4>GFP |  |  | Expression of a cytoplasmic GFP, specifically in *hh* expressing cells |
| **hh-Gal4 (control)**<br>*w<sup>1118</sup>; Gal80<sup>ts</sup>/+; hh-GAL4/UAS: GFP NLS*<br>**<u>Figure 2c-d, 1f-f'', S5t-u</u>** | Gal80<sup>ts</sup> | Gal4>GFP NLS |  |  | Overexpression of GFP NLS (neutral transgene) in hh expressing cells using the UAS Gal4 binary system and the thermosensitive variant of Gal80, repressor of Gal4. |
| **Control**<br>$w^{1118}; UAS:GFP::NLS/ddc-Gal4$<br>**<u>Figure 1g-g'', 1i-j, S2f-f''</u>** | | | Ddc-Gal4>GFP<br>NLS | | Expression of a nuclear GFP, specifically in Dopaminergic neurons |
| **$hh^{ts}/hh^{ts}$**<br>$w^{1118}; UAS:GFP::NLS, hh^{ts2}/ddc-Gal4, hh^{ts2}$<br>**<u>Figure 1h-h'', 1i-1j, S2g-g''</u>** | $hh^{ts}/hh^{ts};$ | | Ddc-Gal4>GFP<br>NLS | | Expression of a nuclear GFP, specifically in Dopaminergic neurons in a $hh^{ts2}/hh^{ts2}$ mutant background. |
| **mCD8::GFP/ddc-Gal4**<br>$w^{1118}; UAS:mCD8::GFP/ddc-Gal4$<br>**<u>Figure 1k-l, 1o S2a-b, S2e</u>** | | | Ddc-Gal4>mCD8::GFP | | Expression of a membrane bound mCD8::GFP, specifically in Dopaminergic neurons |
| **$hh^{ts}/hh^{ts};$**<br>**mCD8::GFP/ddc-Gal4**<br>$w^{1118}; UAS:mCD8::GFP, hh^{ts2}/ddc-Gal4, hh^{ts2}$<br>**<u>Figure 1m-n, 1o, S2c-d, S2e</u>** | $hh^{ts}/hh^{ts};$ | | Ddc-Gal4>mCD8::GFP | | Expression of a membrane bound mCD8::GFP, specifically in Dopaminergic neurons in a $hh^{ts2}/hh^{ts2}$ mutant background. |
| **$w^{1118}$ (control)**<br>$w^{1118}$<br>**<u>Figure 2a-b, 1d-d', S3c-d, S3f-j</u>** | | | | | fly strain with $w^{1118}$ control genetic background |
| **$hh^{ts}/hh^{ts}$ mutant**<br>$w^{1118}; hh^{ts2}/hh^{ts2}$<br>**<u>Figure 2a-b, S3c</u>** | $hh^{ts2}/hh^{ts2}$ | | | | A $hh$ temperature sensitive allele ($hh^{ts2}$), at permissive temperature (18°C) is inactive, at non-permissive temperature (29°), is considered a *hh* null. |
| ***hh<sup>ts</sup>/hh<sup>15</sup>* mutant**<br>*w<sup>1118</sup>; hh<sup>ts2</sup>/hh<sup>15</sup>*<br>**<u>Figure 2a</u>** | *hh<sup>ts2</sup>/hh<sup>15</sup>* |  |  |  | A heteroallelic combination of the *hh* temperature sensitive allele (*hh<sup>ts2</sup>*) and the null allele *hh<sup>15</sup>*. |
| ***hh*-Gal4 (control)**<br>*w<sup>1118</sup>; Gal80<sup>ts</sup> /+; hh-GAL4/UAS:GFP RNAi*<br>**<u>Figure 2a.</u>** | Gal80<sup>ts</sup> | Gal4>GFP RNAi |  |  | Overexpression of GFP RNAi (neutral RNAi) in *hh* expressing cells using the UAS Gal4 binary system and the thermosensitive variant of Gal80, repressor of Gal4. |
| ***hh* OE (*hh*-Gal4):**<br>*w<sup>1118</sup>; Gal80<sup>ts</sup> /+; hh-Gal4, UAS:HhM4/UAS:GFP RNAi*<br>**<u>Figure 2a-b</u>** | Gal80<sup>ts</sup> | Gal4>Hh M4 |  |  | Overexpression of *hh* in *hh*-expressing cells using the UAS Gal4 binary system and the thermosensitive variant of Gal80, repressor of Gal4. At 29°C, Gal80 activity is abolished, leading to the full activity of Gal4 under *hh* promoter |
| ***hh* OE (*hh*-Gal4):**<br>*w<sup>1118</sup>; Gal80<sup>ts</sup>/+; hh*<br>*Gal4,UAS:HhM4/UAS:GFP*<br>*NLS*<br>**<u>Figure 2c-d, S5t-u</u>** | Gal80<sup>ts</sup> | Gal4>Hh<br>M4 |  |  | See above -<br>*hh* OE (*hh*-<br>Gal4):<br>explanation |
| ***hh* OE + *elav*-Gal80:**<br>*w<sup>1118</sup>; Gal80<sup>ts</sup>/elav-Gal80;*<br>*hh-GAL4,UAS:HhM4/+*<br>**<u>Figure 2c-d</u>** | Gal80<sup>ts</sup> | Gal4>Hh<br>M4 | Gal80 |  | Hh OE with<br>constitutive<br>expression of<br>the Gal80<br>repressor<br>which<br>represses<br>expression of<br>Gal4 and thus<br>the UAS Hh<br>transgene in all<br>Elav positive<br>neurons. |
| ***hh* OE + *repo*-Gal80:**<br>*w<sup>1118</sup>; Gal80<sup>ts</sup>/repo-Gal80;*<br>*hh-GAL4,UAS:HhM4/+*<br>**<u>Table S2</u>** | Gal80<sup>ts</sup> | Gal4>Hh<br>M4 |  | Gal80 | Hh OE with<br>constitutive<br>expression of<br>the Gal80<br>repressor<br>which<br>represses<br>expression of<br>Gal4 and thus<br>the UAS Hh<br>transgene in all<br>Repo positive<br>glia. |
| ***hh* mutant + *hh* OE (*vGlut*-**<br>**Gal4)**<br>*w<sup>1118</sup>; Gal80<sup>ts</sup>;UAS: HhM4,*<br>*hh<sup>ts2</sup>/vGlut-GAL4, hh<sup>ts2</sup>*<br>**<u>Figure 2e-f</u>** | *Gal80<sup>ts</sup> ;*<br>*hh<sup>ts2</sup>/hh<sup>ts2</sup>* |  | vGlut-Gal4>HhM4 |  | Overexpression<br>of *hh* in<br>glutamatergic<br>neurons in a<br>*hh<sup>ts2</sup>/hh<sup>ts2</sup>*<br>mutant<br>background. |
| ***hh* mutant (*vGlut*-Gal4)**<br>*w<sup>1118</sup>; Gal80<sup>ts</sup>;UAS:GFP*<br>*NLS, hh<sup>ts2</sup>/vGlut-GAL4, hh<sup>ts2</sup>*<br>**<u>Figure 2e-f</u>** | *Gal80<sup>ts</sup> ;*<br>*hh<sup>ts2</sup>/hh<sup>ts2</sup>* |  | vGlut Gal4>GFP<br>NLS |  | Overexpression<br>of GFP-NLS in<br>glutamatergic<br>neurons in a<br>*hh<sup>ts2</sup>/hh<sup>ts2</sup>* |
|  |  |  |  |  | mutant background. |
| **vGlut-Gal4 (control)**<br>*w<sup>1118</sup>; Gal80<sup>ts</sup>; UAS:GFP NLS/vGlut-GAL4*<br>**Figure 2e-f** | *Gal80<sup>ts</sup>;* |  | vGlut Gal4>GFP NLS |  | Overexpression of GFP NLS in glutamatergic neurons |
| **hh mutant + hh OE (Gad-Gal4)**<br>*w<sup>1118</sup>; Gal80<sup>ts</sup>; UAS: HhM4, hh<sup>ts2</sup>/Gad-Gal4, hh<sup>ts2</sup>*<br>**Figure S3a-b** | *Gal80<sup>ts</sup>;*<br>*hh<sup>ts2</sup>/hh<sup>ts2</sup>* |  | Gad Gal4>HhM4 |  | Overexpression of *hh* in GABAergic neurons in a *hh<sup>ts2</sup>/hh<sup>ts2</sup>* mutant background. |
| **hh mutant (Gad-Gal4)**<br>*w<sup>1118</sup>; Gal80<sup>ts</sup>; UAS:GFP NLS, hh<sup>ts2</sup>/Gad-Gal4, hh<sup>ts2</sup>*<br>**Figure S3a-b** | *Gal80<sup>ts</sup>;*<br>*hh<sup>ts2</sup>/hh<sup>ts2</sup>* |  | Gad Gal4>GFP NLS |  | Overexpression of GFP NLS in GABAergic neurons in a *hh<sup>ts2</sup>/hh<sup>ts2</sup>* mutant background. |
| **Gad-Gal4 (control)**<br>*w<sup>1118</sup>; Gal80<sup>ts</sup>; UAS:GFP-NLS/Gad-Gal4*<br>**Figure S3a-b** | *Gal80<sup>ts</sup>;* |  | Gad Gal4>GFP NLS |  | Overexpression of GFP NLS in GABAergic neurons |
| **RNAi *ci* (repo-Gal4)**<br>*w<sup>1118</sup>; UAS: Dcr2 /Gal80<sup>ts</sup>; UAS: Ci RNAi /Repo-Gal4*<br>**Figure 3a, S1e-e"** | *Gal80<sup>ts</sup>;* |  |  | Repo-Gal4> RNAi *ci* | Glial-specific RNAi knockdown of *ci* |
| **RNAi *smo* (repo-Gal4)**<br>*w<sup>1118</sup>; UAS: smo RNAi/+; smo<sup>X42</sup>, UAS: Dcr2/Gal80<sup>ts</sup>; Repo-GAL4/+*<br>**Figure 3a, S1c-c"** | *Gal80<sup>ts</sup>;*<br>*smo<sup>X42</sup>* |  |  | Repo-Gal4> RNAi *smo* | Glial-specific RNAi knockdown of *smo* in a *smo<sup>X42</sup>* heterozygote mutant background. |
| **control (repo-Gal4)** | *Gal80<sup>ts</sup>;* |  |  | Repo-Gal4> RNAi GFP | Glial-specific knockdown of |
| *w<sup>1118</sup>*; UAS:GFP RNAi/<br>*Gal80<sup>ts</sup>*; UAS:Dcr2/Repo-GAL4<br>**Figure 3a** |  |  |  |  | control GFP<br>RNAi |
| **RNAi *ci* (elav-Gal4)**<br>*w<sup>1118</sup>*, *elav*-Gal4/+; UAS:<br>Dcr2/*Gal80<sup>ts</sup>*; UAS:*ci* RNAi/+<br>**Figure 3b** | *Gal80<sup>ts</sup>*; |  | Elav-Gal4><br>RNAi *ci* |  | Neuron-specific<br>RNAi<br>knockdown of<br>*ci* |
| **RNAi *smo* (elav-Gal4)**<br>*w<sup>1118</sup>*, UAS: *smo* RNAi/elav<br>Gal4; *smo<sup>X42</sup>*,<br>UAS:Dcr2/*Gal80<sup>ts</sup>*<br>**Figure 3b** | *Gal80<sup>ts</sup>*;<br>*smo<sup>X42</sup>* |  | Elav-Gal4><br>RNAi *smo* |  | Neuron-specific<br>RNAi<br>knockdown of<br>*smo* in a<br>*smo<sup>X42</sup>*<br>heterozygote<br>mutant<br>background. |
| **control (elav-Gal4)**<br>*w<sup>1118</sup>*, *elav*-<br>Gal4/+; UAS:Dcr2 UAS:GFP<br>RNAi/*Gal80<sup>ts</sup>*<br>**Figure 3b** | *Gal80<sup>ts</sup>*; |  | Elav-Gal4><br>RNAi GFP |  | Neuron-specific<br>knockdown of<br>control GFP<br>RNAi |
| ***ci<sup>act</sup>* OE (repo-Gal4) in *hh***<br>**mutant**<br>*w<sup>1118</sup>*; *Gal80<sup>ts</sup>*; UAS:*ci<sup>act</sup>*,<br>*hh<sup>ts2</sup>*/repo-Gal4, *hh<sup>ts2</sup>*<br>**Figure 3c-d, 3i, 3l-o, 4b**<br>**S3k S4a-a”** | *Gal80<sup>ts</sup>*;<br>*hh<sup>ts2</sup>*/*hh<sup>ts2</sup>* |  |  | Repo-Gal4> *ci<sup>act</sup>* | Overexpression<br>of an activated<br>form of *ci* (*ci<sup>act</sup>*)<br>in glia in<br>*hh<sup>ts2</sup>*/*hh<sup>ts2</sup>*<br>mutant<br>background. |
| ***hh* mutant (repo-Gal4)**<br>*w<sup>1118</sup>*; *Gal80<sup>ts</sup>*; UAS:GFP<br>NLS, *hh<sup>ts2</sup>*/repo-Gal4, *hh<sup>ts2</sup>*<br>**Figure 3c-d, 3h, 3k, 3m-o,**<br>**4b, 4e-f, 4h, 4l-p S3k S4a-**<br>**d”, S5S** | *Gal80<sup>ts</sup>*;<br>*hh<sup>ts2</sup>*/*hh<sup>ts2</sup>* |  |  | Repo-Gal4> GFP<br>NLS | Overexpression<br>of GFP-NLS in<br>glia in<br>*hh<sup>ts2</sup>*/*hh<sup>ts2</sup>*<br>mutant<br>background. |
| **control (repo-Gal4)**<br>*w<sup>1118</sup>*; UAS:GFP NLS/ *Gal80<sup>ts</sup>*;<br>Repo-GAL4 | *Gal80<sup>ts</sup>*; |  |  | Repo-Gal4> GFP<br>NLS | Overexpression<br>of GFP-NLS in<br>glia |
| **<u>Figure 3c-g, 3j, 3m-o, 4b, 4g-g', 4i 4l-p, 5a-c, 5e-e'', 5i-m, S3k S4a-d'', S5r-s, S6b-b'', S6d-d'', S6f-f'', S6h-h''</u>** |  |  |  |  |  |
| ***ci*<sup>act</sup> OE (repo-Gal4)**<br>*w*<sup>1118</sup>, *Gal80*<sup>ts</sup>, *UAS:ci*<sup>act</sup>/*repo-Gal4*<br>**<u>Figure 3e</u>** | *Gal80*<sup>ts</sup>; |  |  | Repo-Gal4> *ci*<sup>act</sup> | Overexpression of an activated form of *ci* (*ci*<sup>act</sup>) in glia |
| ***smo*<sup>act</sup> OE (repo-Gal4)**<br>*w*<sup>1118</sup>, *Gal80*<sup>ts</sup>, *UAS:smo*<sup>act</sup>/*repo-Gal4*<br>**<u>Figure 3f</u>** | *Gal80*<sup>ts</sup>; |  |  | Repo-Gal4> *smo*<sup>act</sup> | Overexpression of an activated form of *smo* (*smo*<sup>act</sup>) in glia |
| **16xLexAop2-mCD8GFP/5x mCD8-Cherry; *ci*-LexA/*repo-Gal4***<br>*w*<sup>1118</sup>, 16xLexAop2-mCD8GFP/5xmCD8-Cherry; *ci*-LexA/*repo-Gal4*<br>**<u>Figure S1a-a''</u>** |  |  |  | Repo-Gal4>mCD8::Cherry | Expression of a membrane-bound mCD8::Cherry, specifically in glia (*repo-Gal4*) along with expression of mCD8GFP in Ci expressing cells (using the lexAop-lexOp system) |
| ***hsp40* OE (repo-Gal4) in *hh* mutant**<br>*w*<sup>1118</sup>, *Gal80*<sup>ts</sup>, *UAS:hsp40*, *hh*<sup>ts2</sup>/*repo-Gal4*, *hh*<sup>ts2</sup><br>**<u>Figure 4j, 4l-l', S4c-c''</u>** | *Gal80*<sup>ts</sup>;<br>*hh*<sup>ts2</sup>/*hh*<sup>ts2</sup> |  |  | Repo-Gal4> *hsp40* | Overexpression of *hsp40* in glia in *hh*<sup>ts2</sup>/*hh*<sup>ts2</sup> mutant background. |
| ***hsp68* OE (repo-Gal4) in *hh* mutant**<br>*w*<sup>1118</sup>, *Gal80*<sup>ts</sup>, *UAS:hsp68*, *hh*<sup>ts2</sup>/*repo-Gal4*, *hh*<sup>ts2</sup><br>**<u>Figure 4k-p, S4b-b''</u>** | *Gal80*<sup>ts</sup>;<br>*hh*<sup>ts2</sup>/*hh*<sup>ts2</sup> |  |  | Repo-Gal4> *hsp68* | Overexpression of *hsp68* in glia in *hh*<sup>ts2</sup>/*hh*<sup>ts2</sup> mutant background. |
| ***hsp23* OE (repo-Gal4) in *hh* mutant** | *Gal80*<sup>ts</sup>;<br>*hh*<sup>ts2</sup>/*hh*<sup>ts2</sup> |  |  | Repo-Gal4> *hsp23* | Overexpression of *hsp23* in glia |
| $w^{1118}; Gal80^{ts}; UAS:hsp23, hh^{ts2}/repo-Gal4, hh^{ts2}$<br>**Figure S4d-d''** | | | | | in $hh^{ts2}/hh^{ts2}$ mutant background. |
| $hh^{ts}/hh^{ts}; mCD8::GFP; repo-Gal4$<br>$w^{1118}; UAS:mCD8::GFP, hh^{ts2}/repo-Gal4, hh^{ts2}$<br>**Figure S5e-f''; S5i-j''** | $hh^{ts2}/hh^{ts2}$ | | | Repo-Gal4>mCD8::GFP | Expression of a membranous bound mCD8::GFP, specifically in glia in a $hh^{ts2}/hh^{ts2}$ mutant background. |
| **ci-Gal4; mCD8-GFP**<br>$w^{1118}; UAS:mCD8\ GFP; ci-Gal4\ Trojan$<br>**Figure S5o-p''** | | | | | Expression of a membrane bound mCD8::GFP specifically in *ci* expressing cells |
| **Human $A\beta42^{E693G}$ OE; (repo-Gal4) + GFP NLS**<br>$w^{1118}; UAS: A\beta42^{E693G} /UAS: GFP\ NLS; repo-Gal4$<br>**Figure 5a-d''', 5f-f'', 5j-k S6a-a'', c-c'', e-e'', g-g''** | | | | Repo-Gal4><br>$A\beta42^{E693G}$ | Overexpression of human artice variant $A\beta42^{E693G}$ and GFP NLS in glia |
| **Human $A\beta42^{E693G}$ OE + $Smo^{Act}$ OE; (repo-Gal4)**<br>$w^{1118}; UAS: A\beta42^{E693G} /UAS: Smo^{Act}; repo-Gal4$<br>**Figure 5a, 5c, 5g-g'', 5j-k, S6i-i''** | | | | Repo-Gal4><br>$A\beta42^{E693G}/Smo^{Act}$ | Overexpression of human artice variant $A\beta42^{E693G}$ and $Smo^{Act}$ in glia |
| **Human $A\beta42^{E693G}$ OE; (repo-Gal4) + Hsp68**<br>$w^{1118}; UAS: A\beta42^{E693G} ;UAS:Hsp68/ repo-Gal4$<br>**Figure 5b-c, h-h'', 5j-k, S6j-j''** | | | | Repo-Gal4><br>$A\beta42^{E693G};hsp68$ | Overexpression of human artice variant $A\beta42^{E693G}$ and *hsp68* in glia |
| **control (*repo-Gal4*)**<br>*w<sup>1118</sup>*; *UAS:GFP NLS*;<br>*Repo-GAL4*<br>**<u>Figure 5a-c, e-e''', i-k,</u>**<br>**<u>S6b-b''', d-d''', f-f''', h-h'''</u>** |  |  |  | Repo-Gal4> GFP<br>NLS | Overexpressio<br>n of GFP-NLS<br>in glia |

**Table S5.** Shows the data sets that were compared when calculating the p value in the lifespan studies.

| Data Set 1 |  | Control Data Set 2 |  | P value | Figure |
| --- | --- | --- | --- | --- | --- |
| Genotype | Median Survival Time | Genotype | Median Survival Time |  |  |
| *hh* OE (*hh*-Gal4) | 48d | *hh*-Gal4 (control) | 38d | P<0.0001 | 2a |
| *hh<sup>ts</sup>/hh<sup>ts</sup>* mutant | 8d | *w<sup>1118</sup>* (control) | 38d | P<0.0001 | 2a |
| *hh<sup>ts</sup>/hh<sup>15</sup>* mutant | 6d | *w<sup>1118</sup>* (control) | 38d | P<0.0001 | 2a |
| *hh* OE + *elav*-Gal80: | 38d | *hh* OE (*hh*-Gal4) | 43d | P<0.0001 | 2c |
| *hh* OE (*hh*-Gal4) | 43d | *hh*-Gal4 (control) | 37d | P<0.0001 | 2c |
| *hh* mutant + *hh* OE (*vGlut*-Gal4) | 28d | *hh* mutant (*vGlut*-Gal4) | 7d | P<0.0001 | 2e |
| *hh* mutant + *hh* OE (*Gad*-Gal4) | 18d | *hh* mutant (*Gad*-Gal4) | 8d | P<0.0001 | S3a |
| RNAi *smo* (*repo*-Gal4) | 25d | control (*repo*-Gal4) | 44d | P<0.0001 | 3a |
| RNAi *ci* (*repo*-Gal4) | 23d | control (*repo*-Gal4) | 44d | P<0.0001 | 3a |
| *Ci<sup>act</sup>* OE (*repo*-Gal4) in *hh* mutant | 31d | *hh* mutant (*repo*-Gal4) | 6d | P<0.0001 | 3c |
| *hsp40* OE (*repo*-Gal4) in *hh* mutant | 28d | *hh* mutant (*repo*-Gal4) | 7d | P<0.0001 | S4c |
| *hsp68* OE (*repo*-Gal4) in *hh* mutant | 31d | *hh* mutant (*repo*-Gal4) | 6d | P<0.0001 | 4o, S4b'' |
| *ci<sup>act</sup>* OE (*repo*-Gal4) | 48d | control (*repo*-Gal4) | 42d | P<0.0001 | 3e |
| *smo<sup>act</sup>* OE (*repo*-Gal4) | 49d | control (*repo*-Gal4) | 41d | P<0.0001 | 3f |
| Human *Aβ42<sup>E693G</sup>* OE; (*repo*-Gal4) | 22d (20% survival time) | Human *Aβ42<sup>E693G</sup>* OE/ *Smo<sup>Act</sup>* OE;(*repo*-Gal4) | 26d (20% survival time) | P<0.0001 | 5a |
| Human *Aβ42<sup>E693G</sup>* OE; (*repo*-Gal4) | 22d (20% survival time) | Human *Aβ42<sup>E693G</sup>* OE; *hsp68* OE (*repo*-Gal4) | 26d (20% survival time) | P<0.0001 | 5b |

**Table S6.** Displays the data sets that were compared when calculating the p value in the locomotion assays.

| Data Set 1 |  | Control Data Set 2 |  | P value | Figure |
| --- | --- | --- | --- | --- | --- |
| Genotype | Climbing activity (%) | Genotype | Climbing activity (%) |  |  |
| *hh* OE (*hh*-Gal4) | 88% | *w<sup>1118</sup>* (control) | 66% | P<0.001 | 2b |
| *hh<sup>ts</sup>/hh<sup>ts</sup>* mutant | 4% | *w<sup>1118</sup>* (control) | 66% | P<0.001 | 2b |
| *hh* OE + *elav*-Gal80: | 20% | *hh* OE (*hh*-Gal4) | 39% | P<0.001 | 2d |
| *hh* mutant + *hh* OE (*vGlut*-Gal4) | 55% | *hh* mutant (*vGlut*-Gal4) | 26% | P<0.001 | 2f |
| *hh* mutant + *hh* OE (*Gad*-Gal4) | 45% | *hh* mutant (*Gad*-Gal4) | 22% | P<0.001 | S3b |
| *Ci<sup>act</sup>* OE (*repo*-Gal4) in *hh* mutant | 54% | *hh* mutant (*repo*-Gal4) | 17% | P<0.001 | 3d |
| *hsp68* OE (*repo*-Gal4) in *hh* mutant | 38% | *hh* mutant (*repo*-Gal4) | 10% | P<0.001 | 4p |
| Human *Aβ42<sup>E693G</sup>* OE/ *Smo<sup>Act</sup>* OE (*repo*-Gal4) | 56% | Human *Aβ42<sup>E693G</sup>* OE; (*repo*-Gal4) | 21% | P<0.001 | 5c |
| Human *Aβ42<sup>E693G</sup>* OE; *hsp68* OE (*repo*-Gal4) | 66% | Human *Aβ42<sup>E693G</sup>* OE; *hsp68* OE (*repo*-Gal4) | 21% | P<0.001 | 5c |

